# Ballistic Microscopy (BaM)

**DOI:** 10.1101/2025.10.07.681030

**Authors:** A. S. Jijumon, Ray Chang, Manu Prakash

## Abstract

Light and electron microscopy utilizes interactions of either photons or electrons with matter to create images from cellular to atomic scale. However, these methods are limited in de novo discovery and spatial mapping of unknown biomolecules. Label free methods such as mass spectrometry or sequencing lack live-cell and subcellular context. Here we introduce a new approach, Ballistic Microscopy (BaM), to image cells with physical nanoparticles. We bombard living cells with millions of nanoparticles traveling at ∼1000 m/s. Each particle passes through cells, piercing and capturing attoliters of cytoplasm on a hydrogel substrate while preserving spatial information (SPLAT-MAP). This “physical image” of a live cell captures a molecular fingerprint of a cell on a hydrogel film that can be processed post-capture via multiple techniques such as TEM, Cryo-EM, mass spectrometry, confocal imaging, and DNA amplification. Using BaM, we discover previously unknown composition of CLIP170 and Tau3R condensates in HEK cells, uncovering Keratin-18 as a structural element. BaM establishes a new paradigm of “physical imaging” with modular readout platform for spatially resolved live sampling across cells, tissues, and organisms.

---

A living cell is a highly complex, dense, heterogeneous, and dynamic architecture of thousands of biomolecules, including proteins, nucleic acids, lipids, cytoskeletal assemblies, organelles, and more, organized in both space and time [1][2]. While genomics has garnered significant attention, the vast majority of cellular functions in a living cell are executed at level of proteins and protein assemblies [3]. Understanding the spatial organization, interactions, and dynamics of proteins within live cells is thus essential for deciphering both physiological and pathological states of a cell [4]. Despite advances in protein biology [5][6][7], a large fraction of the human proteome remains uncharacterized, often referred to as the “dark proteome,” with unknown three-dimensional structures or any identifiable homologs or specific spatial distribution patterns within cell types [8]. This fraction is even higher in non-model organisms [9]. Compounding the challenge, most proteins have multiple interaction partners, necessitating methods that can probe unknown and unlabeled bio-molecular interactions in their native context [10].

Conventional imaging-based techniques for studying genes, proteins, and metabolites in live cells typically rely on tagging known analytes, such as using fluorescent proteins like GFP [11][12][13]. While powerful, these methods are limited to predefined known targets, leaving a vast portion of cellular content unexplored [14]. In contrast, biochemical approaches such as co-immunoprecipitation [15][16], proximity ligation assays [17][18], and affinity purification mass spectrometry [19][20] provide deeper insights into molecular interactions but are generally performed on fixed or lysed cells and lack spatial resolution. Emerging live-cell subsampling methods, including AFM probes [21][22], nanostraws [23], carbon nanotube spearing [24][25], and microneedle techniques [26], physically puncture cells to extract intracellular material. However, they generally collect the cytoplasm as a bulk, undifferentiated mixture. These approaches often lack subcellular spatial resolution, are technically demanding, have very low throughput, and are typically limited to a few specialized laboratories and cell types.

Here we introduce a new paradigm in microscopy - Ballistic Microscopy (BaM), a label-free method that physically subsamples local nanoscale regions of live cells using high-speed ballistic nanoparticles. Rather than imaging with photons or electrons, BaM samples cytoplasm and membrane components via nano-particle penetration piercing the entire cell and consecutive capture on the other side. This is akin to a ”physical image” being captured on a hydrogel ”film” with physical material from a cell captured on these nano-bullets. These nanoparticles, ranging from 50 to 1000 nanometers in diameter, are accelerated en-masse, traveling as a physical particle beam at velocities nearing one kilometer/sec piercing through a living cell and captured on a stable substrate while maintaining spatial localization. Remarkably, these individual particles collect attoliter-scale volumes of cytoplasm from specific entry locations within the cell. The harvested material mapped onto individual nano-particles for every BaM shot can then be analyzed offline using downstream high-resolution techniques such as mass spectrometry or cryo-electron microscopy. BaM uniquely decouples the step of sampling and detection, enabling a modular framework where a single sample can be subjected to multiple orthogonal readouts, including structural or biochemical analyses. Although ballistics is commonly been used in the past to introduce materials into a cell [27][28], here we flip it’s function and use it to extract spatially organized material from single cells. This approach creates a new paradigm for exploring the molecular composition of unlabeled subcellular environments and assemblies in situ creating “physical images” of living cells.

We demonstrate the utility of BaM by extracting cytosolic and membrane material from live human HEK293 cells and a non-model protist, Chaos (Pelomyxa) carolinensis. The BaM SPLAT-MAP samples are characterized via numerous techniques including fluorescence microscopy, mass spectrometry, transmission electron microscopy, and cryogenic electron tomography. To validate BaM’s biological utility, we focused on five bait proteins: Actin, CLIP170, Tau-3R, ATIP3, and SALS. These include common cytoskeletal component and proteins known to form condensates or aggregates. BaM enabled the in situ retrieval of protein condensates from live cells for the first time. Mass spectrometry of CLIP170 condensate-enriched BaM particles revealed approximately 641 proteins, including Keratin-18, an unexpected structural organizer inside these condensate droplets, also found in Tau-3R condensates with distinct double emulsion architecture.

This label-free approach offers new access to the molecular composition and architecture of multiprotein structures, including condensates, organelles, and other undiscovered subcellular assemblies, across a wide variety of living cells and tissues. Because a single cell can accommodate thousands of nanoparticles simultaneously, BaM provides high spatial resolution, allowing material to be collected and compared from distinct subcellular regions. BaM enables a new mode of imaging by decoupling sample capture and analysis - akin to an old school ”film” camera - where the physical image can be probed with a number of existent destructive detection methods.

## Ballistic Microscopy concept

Ballistic microscopy (BaM) enables analysis of unlabeled cellular content from live cells using high-speed nanoparticle penetration and capture. Accelerated gold nanoparticles traveling upto ∼1000 m/s pierce individual cells, traversing subcellular distances of ∼2–4 *µ*m for a HEK-cell with an impact duration in the picosecond range (∼ 5 × 10^−12^s) (Fig.1). Each particle extracts attoliter volumes of cytoplasmic material through momentum transfer (Fig. 1A), and slowing down in the process after passing through the cell. The collected samples, embedded within a capture gel film, retain spatial information and are suitable for downstream ‘omics’ studies (Fig. 1B). The exceptionally fast speed of travel enables a clean trajectory and precise capture of atto-liters of cytoplasmic droplets that cling to the surface of the nanoparticle. This process is analogous to a high-speed bullet extracting trace materials while maintaining sufficient momentum to exit. By integrating downstream high-resolution structural and compositional analysis from the same sample, BaM offers a powerful platform in the future for studying live cells at unprecedented spatio-temporal resolution (Fig. 1B).

**Figure 1:**
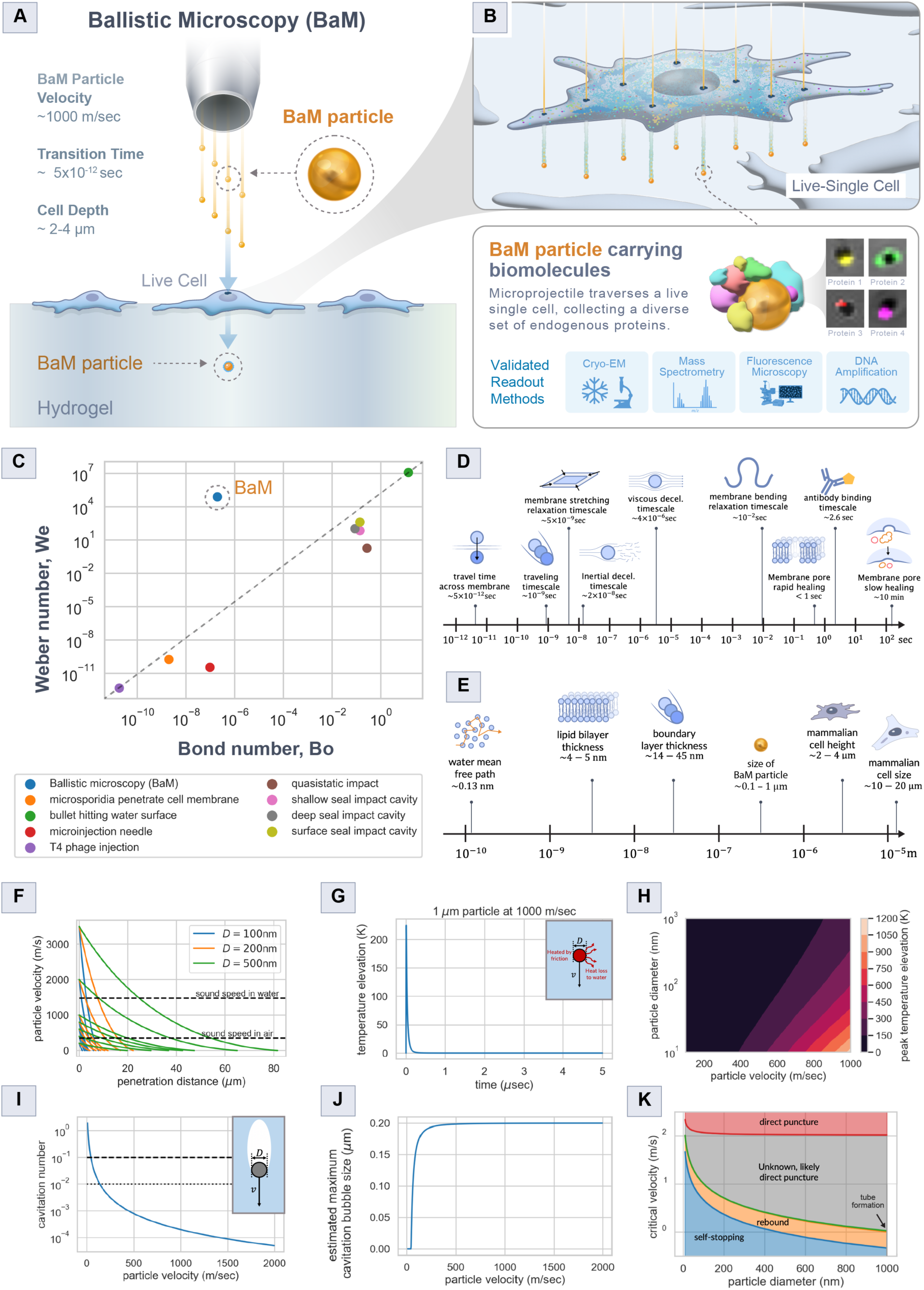
Ballistic Microscopy concept and theoretical assessment of BaM. (A) Schematic of the BaM concept. High-speed nanoparticles launched toward live cells on a particle-penetrable substrate. The impact duration is in the picosecond range (∼ 5 × 10^−12^ s), allowing particles to traverse a subcellular distance (∼2–4 *µ*m) and collect cellular materials. (B) Conceptual diagram illustrating cytoplasm subsampling by BaM and tagged proteins captured on individual BaM particles (pseudo-colored). BaM is compatible with multiple orthogonal downstream readouts, including Cryo-EM, mass spectrometry, fluorescence microscopy, and DNA amplification.(C) Dimensionless phase diagram shows that BaM occupies a unique regime of high Weber and low Bond number, distinct from other high-speed interfacial phenomena. (D&E) BaM operates under extreme separations of time scale (D) and length scale (E): a 1 *µ*m gold particle traveling at 1000 m/s crosses the membrane in picoseconds, minimizing surface chemistry effects during sampling. (F) Particles ≥ 200 nm can traverse typical cell heights (2-4 *µ*m) at ≥ 200 m/s, while smaller particles require higher velocities. (G&H) Transient thermal modeling of a 1 *µ*m gold particle traveling at 1000 m/s shows a 200K peak temperature rise that dissipates within 0.5 *µ*s; larger particles experience lower heating. (I&J) Cavitation is likely during BaM with cavitation number (Cav) <0.1; bubble growth remains at submicron scale due to short transit time. (K) Particle-membrane interaction regimes show efficient sampling occurs in the direct puncture regime, with hydrodynamic drag suppressing tube formation.

## Theoretical considerations

The design of this unusual instrument requires unique theoretical considerations. For this we studied the unique physical regime that BaM exists in - defined by very small particle size (sub-micron) and extreme velocity (∼1000 m/s). Based on the typical parameter values associated with the process (length scale *L* ∼ particle diameter *D* ∼ 1*µ*m, fluid density *p*_f_ ∼ 1000 kg/m^3^, interfacial tension *σ* ∼ 50 mN/m[29], velocity scale *U*_0_ ∼ 1000 m/s, dynamic viscosity *µ* ∼ 1 mPa-s, speed of sound *c* ∼ 1500 m/s, gravitational acceleration *g* ∼ 9.8 m/s^2^), we find that BaM is a rare example of a low-Bond-number, high-Weber-number, high-Reynolds-number, high-capillary-number, and intermediate-to-high-Mach-number interfacial phenomenon (Table S1).

Compared to other macroscopic high-speed penetrating interfacial phenomena (Fig. 1C), BaM falls into a unique region of the Weber-Bond number phase space. While most other penetrating phenomena lie along the diagonal, BaM is distinctly off-diagonal in a low-Bond-number, high-Weber-number quadrant. We can understand this intuitively by looking at the ratio between Weber number and Bond number. As the interfacial tension term cancel out with each other, the ratio of the two becomes We/Bo = *U*^2^/(*gL*) ∼ (*aL*)/(*gL*) ∼ *a*/*g*, where *a* is the characteristic acceleration assuming the particle is accelerated for a length comparable to its own length. The fact that microparticles in BaM is off-diagonal from other penetrating interfacial phenomena indicates its extraordinarily high acceleration in its mechanism.

BaM also involves drastic separations in both time and spatial scales (Figs. 1D-E). For example, a1 *µ*m particle can traverse a 5-nm lipid bilayer in about 5 ps, which is orders of magnitude shorter than any mechanical or biochemical membrane relaxation, such as bending or protein binding (Fig. 1D). These differences in timescale suggest that cellular response mechanisms, such as membrane repair or protein binding, are too slow to interfere with sampling during BaM. The spatial scales also span several orders of magnitude (Fig. 1E), ranging from molecular free paths and bilayer thickness to whole-cell dimensions. This vast length-scale separation presents challenges for simulation of this high-speed phenomena. In the following, we thus adopt a scaling approach to gain intuitions on the design space available for effective BaM approaches.

Successful sampling of cytoplasm requires the particle to traverse the thickness of the cell without causing damage from high temperature and/or cavitation, and to penetrate the bottom membrane of the cell to escape and be captured by the hydrogel film. Using Brown and Lawler’s correlation [30], we estimate the drag on a speeding nano-particle traversing the cytoplasm (Eq. S1) and estimate the penetration distance of these particles as a function of impingement velocity (see Supplementary text for details). As shown in Figure 1F, particles with diameters ≥200 nm and velocities ≥200 m/s can easily traverse typical mammalian cell heights (∼2–4 *µ*m), assuming cytoplasmic viscosity similar to water. Smaller particles (∼100 nm) require higher velocities (*>*1000 m/s), commonly achieved via standard apparatus.

We next consider whether the penetrating projectile can damage the cell through excessive heating or cavitation. Coupled drag-interfacial thermal conductance equations (Eq. S2) reveal that a1 *µ*m gold particle launched at 1000 m/s may experience a peak temperature increase of ∼200 K, which dissipates within 0.5 *µ*s, much shorter than the typical time scale of thermal denaturation (Fig. 1G). The phase diagram shows that the peak temperature increase is inversely related to particle diameter due to the increased heat capacity and surface area for interfacial thermal conductance (Fig. 1H). These results suggest minimal risk of thermal damage. On the other hand, calculation of the cavitation number for the BaM process (Eq. S3) indicates that cavitation could likely occur, independent of particle size (Fig. 1I). However, because of the rapid transit time, for a 1 *µ*m gold particle, the cavitation bubble cannot grow beyond 0.2 *µ*m (Fig. 1J) (see Supplementary text for details).

To extract cytoplasm, the particle must also penetrate the bottom membrane. Prior MD simulations [31] identified three regimes: rebound, tube formation, and direct puncture. However, previous models ignored hydrodynamic drag. Our re-analysis, incorporating fluid resistance, reveals an additional “self-stopping” regime, where drag alone prevents penetration. This can be visualized as a phase diagram between particle size and critical impinge velocity. We further find that tube formation collapses into an impractically narrow region once drag is considered. Thus, successful sampling must occur in the direct penetration regime (Fig. 1K). An unknown regime is marked on the diagram due to the uncertainty in the thermal velocity of phospholipid on the lipid bilayer membrane (see Supplementary text for details).

Finally, successful sampling requires fragments of the cytoplasm to travel together with the projectile away from the target cells. This implies that the extracted cytoplasm must lie within the boundary layer of the projectile, where momentum transfer occurs. We thus estimated the extracted volume using boundary layer theory. Considering the time over which the fluid experiences acceleration from the particle as τ = *D/υ*, the boundary layer thickness (δ) around the particle can be approximated as 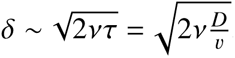, where *a* is the kinematic viscosity of the cytoplasm. Based on this, we can obtain a naive estimation of the volume extracted as *V_ext_* ∼ *πD*^2^*δ* a *D*^5/2^ν^1/2^υ^−1/2^. For a 1 *µ*m particle at 500 m/s, this layer is ∼63 nm thick, corresponding to a sampled volume of 0.2 *µ*m^3^, or 200 aL. Using known protein and mRNA concentrations in mammalian cells [32, 33, 34, 35], this volume likely contains 2 × 10^5^ proteins and ∼20 copies of a specific protein, as well as 0.5–5 mRNA transcripts. These estimates support the feasibility of BaM achieving

single-transcript molecular resolution.

## Design, construction and characterization of BaM

To experimentally test the concept of collecting cellular content from live cells using penetrating ballistic nano-particles, we constructed an in-house BaM setup (Fig. 2A; Fig. S1A-B) consisting of a particle acceleration unit and an imaging unit. Nano-particle acceleration unit was designed to generate high-speed particles which can be accomplished via various previously described mechanisms including compressed gas or light induced shock waves [27][36][37]. In this setup, we used a very well characterized compressed gas approach [38][39][40] (Fig. 2A), similar in architecture to gene-gun based biolistic delivery method [27]. The imaging unit was a modular inverted epi-fluorescence/ brightfield microscope [41] for visualizing fluorescently tagged proteins collected by BaM particles, coupled with an orthogonal (90 degree to imaging axis) high-speed camera (Fig. 1A, 1E; Fig. S1F–G) to capture ultrafast penetration events in real time (Supplementary Video 1).

**Figure 2:**
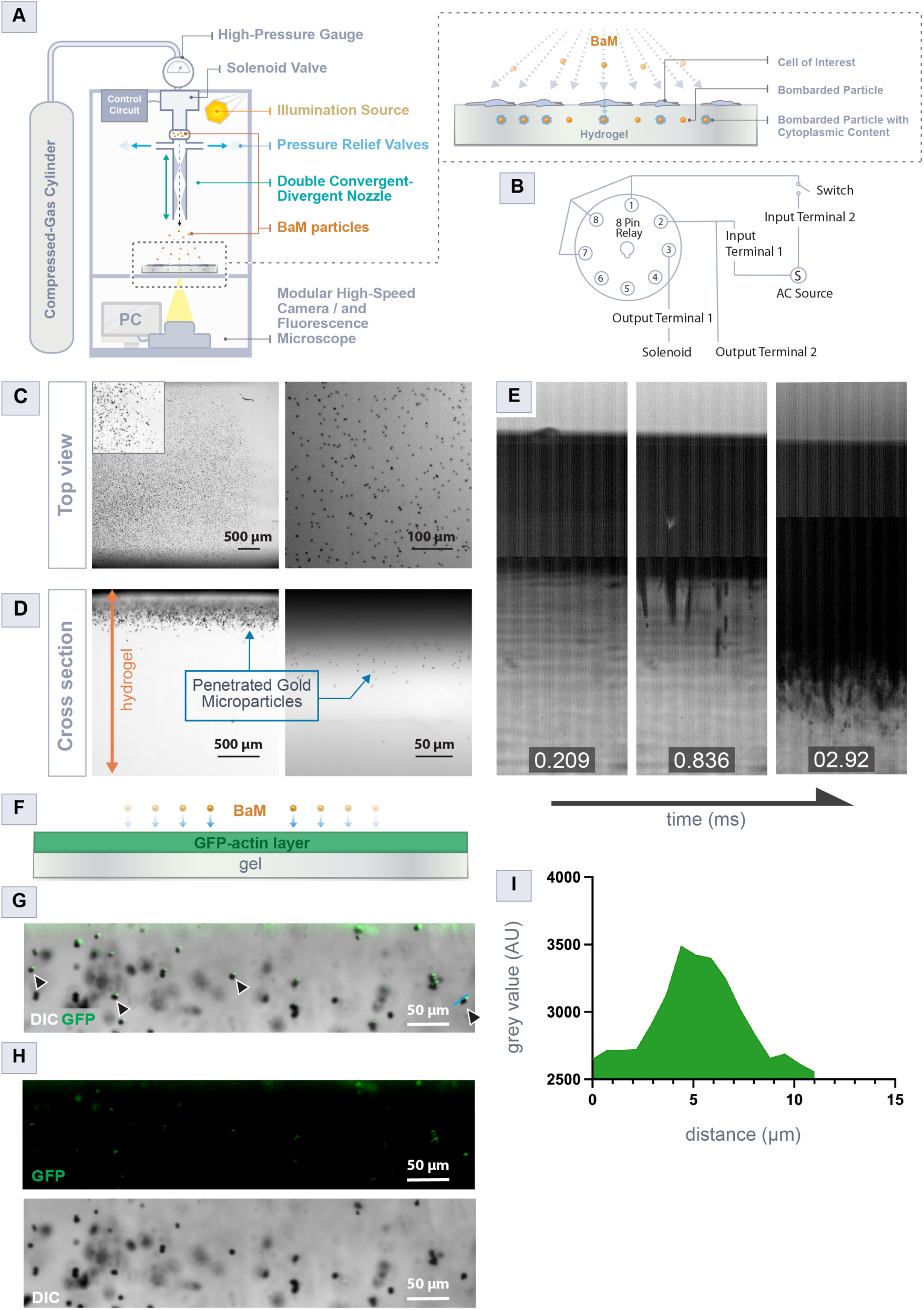
Ballistic Microscopy instrumentation and testing of the BaM concept. (A) Schematic of the BaM setup, which uses a compressed gas-driven particle acceleration. The setup is integrated with a modular fluorescence microscope and a high-speed camera (details in the Methods section). (B) Electrical triggering circuit used to control the solenoid valve via an 8-pin relay. (C–D) Top and cross-sectional views showing gold BaM particles embedded in a gel after bombardment, demonstrating their spatial distribution and penetration depth. (E) High-speed imaging captures the trajectory of particles through the gel over time (see Supplementary Video 1). (F–H) Testing the BaM concept in a non-living system. Clarified cell lysate expressing GFP-tagged actin was coated onto a 1% agarose gel and performed BaM from the top, as illustrated in (F). The side view of a gel cross-section (prepared as shown in Fig. S1e; see Methods) shows merged DIC and fluorescence images, revealing colocalization of the actin-GFP signal (arrowheads) with the penetrated BaM particles (5.1 *µ*m in diameter) (H–I), confirming molecular pickup. n = 3. (I) Quantification of the GFP signal on a representative BaM particle, confirming biomolecule transfer upon impact.

After assembling the BaM setup, we conducted initial tests to characterize depth of particle penetration by firing ballistic particles onto a blank 1 % agarose gels (Fig. 2C). Top view and cross-sectional (Fig. S1 E, also see methods) imaging of the gels with penetrated BaM particles demonstrated that the BaM setup reliably achieved sufficient penetration depths from 100 to 600 *µ* m (Fig. 2D; Fig. S2A-D).

To further characterize the system, we varied key parameters including particle size (0.6 *µ*m , 1 *µ*m , 1.5 *µ*m ), gel concentration (0.5%, 1%, 2%), and gas pressure (50-200 psi) and measured the resulting penetration depths (Fig. S2A-D). These experiments showed that the BaM setup reliably propelled high-speed particles, with penetration scaling monotonically with experimental parameters. Strikingly, using 100-150 psi gas pressure and 0.6-1 *µ*m particle diameters achieved an average penetration depth of 60 *µ*m (Fig. S2D), sufficient to traverse single mammalian cells, which are typically 5-10 *µ*m thick [42]. The setup also achieved penetration depths up to 600 *µ*m with 5.1 *µ*m particles (Fig. S2D), highlighting its potential for sampling thicker tissue specimens, such as brain slices or whole organisms. Using a high-speed camera (Fig. S1F; see Methods), we captured slow-motion video of BaM particles penetrating a gel (Fig. 2E; Fig. S1G; Supplementary Video 1).

After building the BaM setup and optimizing conditions, we tested material collection by high-speed particles in a cell-mimicking system (Fig. 2G-H, Fig. S2E-F). A gel was coated with GFP-tagged actin protein in lysate [43] and impacted from above with BaM particles (see Methods). Following penetration, the gel was cross-sectioned (Fig. S1E) and examined for GFP signals on particles. As a control, BaM particles were also shot onto a gel without any GFP-actin coating (Fig. S2E). Fluorescence microscopy and line-scan analysis confirmed that the high-speed particles penetrating the GFP-actin layer successfully collected GFP, while the GFP-actin coating also reduced penetration depth compared to the control demonstrating increased drag in lysate-coated gels.

In addition to protein collection, we tested BaM particles for nucleic acid capture and amplification (Fig. S2H). Agarose gels coated with a loop-mediated isothermal amplification (LAMP) mix were bombarded with particles, incubated at 65 °C, and screened for fluorescence (also see methods). Merged DIC and fluorescence images confirmed DNA amplification on individual particles, with non-fluorescent particles serving as internal controls. These results demonstrate BaM’s ability to collect both proteins and nucleic acids in a non-living system.

## Isolation of proteins and membrane components from live cells using BaM

After validating BaM on cellular lysate, we next tested its ability to sample biomolecules from live cells. For this, we selected HEK293 adherent cells. We first established a strategy to attach them by placing an electron microscopy (EM) grid on a hydrogel substrate within a glass-bottom dish (Fig. 3A, see Methods). Live HEK293 cells cultured on the grids were stained for nuclei and cell membrane before BaM. This configuration permitted direct spatial correlation between pre-BaM cellular architecture and post-BaM particle-associated fluorescence (Fig. 3B-I; Fig. S3AD). Following BaM and removal of the EM grid, DIC imaging revealed a stencil-like imprint of the grid on the hydrogel, composed of embedded BaM particles (Fig. 3C, 3D, 3F, 3H). The grid like SPLAT-MAP also provides a reference map to original cellular distribution on an EM grid. Fluorescence confocal microscopy detected nuclear and membrane signals on individual nanoparticles, confirming successful material capture during penetration (Fig. 3E, Fig. 3I). Spatial comparison of an EM grid hole before (Fig. 3E; Fig. S3B) and after BaM (Fig. 3F-G; Fig. S3C-D) demonstrated that BaM particles preserved the spatial correspondence to the overlying live cells. Strikingly, Z-stack imaging revealed fluorescence trails along BaM particles, directly mapping their penetration trajectories within the gel (Fig. 3I; Supplementary Video 2).

**Figure 3:**
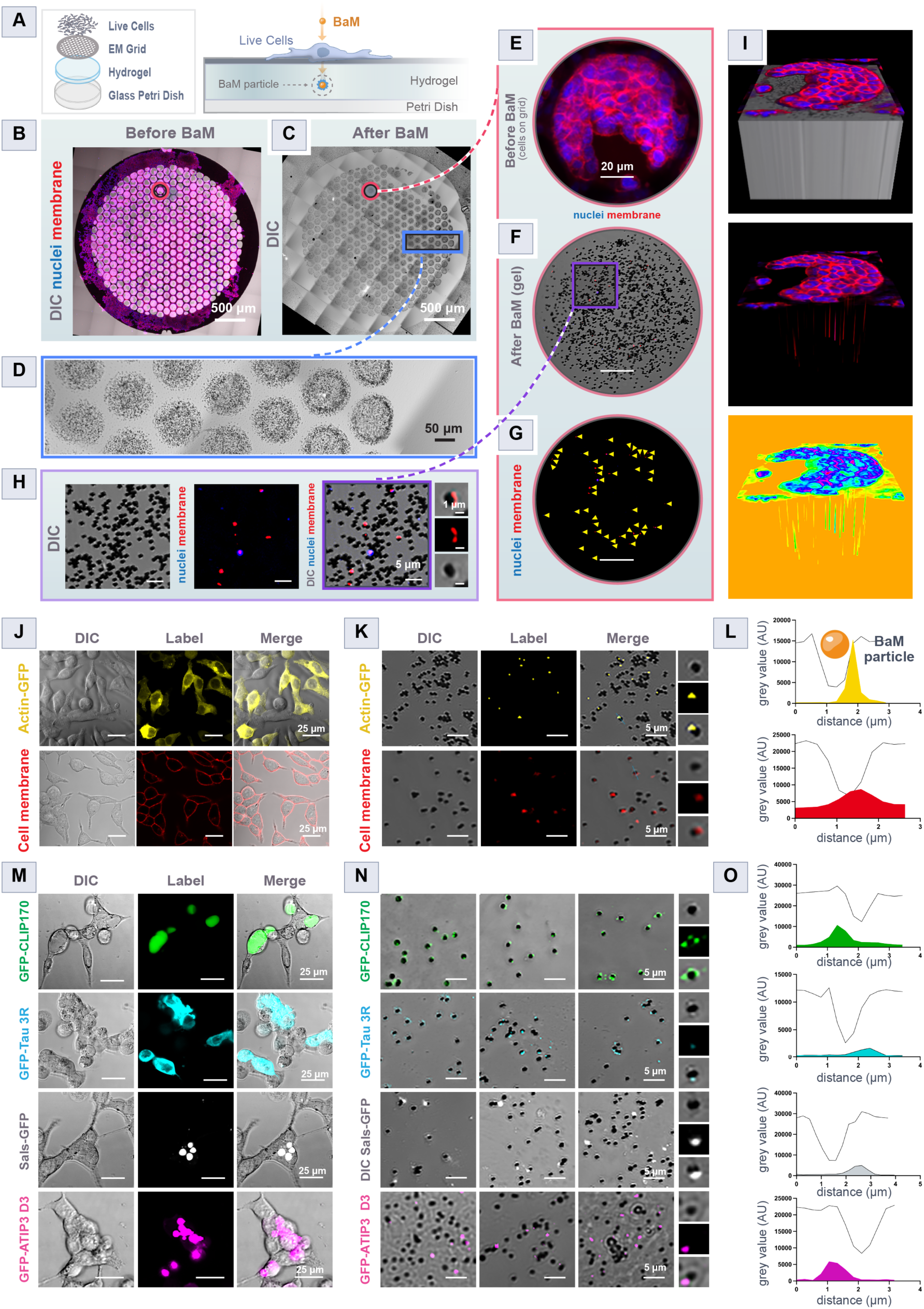
Testing the BaM concept for collecting cytoplasm and membrane from live cells. (A-H) BaM demonstrating the collection of cellular material from live HEK293 cells while retaining spatial information. (A) Schematic of the method used to immobilize cells for BaM. HEK293 cells are cultured on an EM grid, which is then placed onto a hydrogel before BaM. (B-C) DIC and fluorescence images of HEK293 cells stained for nuclei (blue) and membranes (red) before BaM (B), and a DIC image of the gel with embedded BaM particles after BaM (C). (D) Magnified DIC image (blue box in C) showing EM grid holes filled with BaM particles. (E-G) Magnified DIC and fluorescence images of a single EM grid hole shown before and after BaM. (E) Fluorescence image of cells prior to BaM (magenta circle in B), (F) post-BaM (magenta circle in C) showing BaM particles embedded in the gel, and (G) fluorescence signal collected on particles (yellow arrowheads) corresponding to nuclear and membrane markers. (H) Cropped field from (F), showing co-labeled BaM particles with fluorescent signals. Insets highlight individual particles. n = 7-10. (I) 3D reconstruction of an EM grid hole region post-BaM, generated from a Z-stack, showing BaM particles with trailing fluorescence signals. The bottom panel shows false-colored trails for better visualization. (J-0) Validation of the BaM concept on cells cultured directly on a 3D hydrogel. Live HEK293 cells are either stained with live-cell markers or express fluorophore-tagged proteins. (J-K) Cells before BaM, and BaM particles collecting actin-GFP signals (top panel) and membrane signals (bottom panel). Insets show an individual BaM particle. n = 3-5. (L) Line intensity plots for individual BaM particles from (K). (M-O) Application of BaM to live cells expressing various GFP-tagged cytoskeletal condensate- or aggregate- forming proteins: GFP-CLIP170 (green), GFP-Tau 3R (cyan), SALS-GFP (white), and GFP-ATIP3 domain 3 (magenta). Pre-BaM images (M) show fluorescence expressions, while post-BaM gel images (N) display BaM particles carrying GFP signals. n = 3. (O) Line scans of representative BaM particles (from N) confirming fluorescence signals.

Additionally, we successfully applied BaM to cells cultured directly on a 3D hydrogel without the use of an EM grid. In this approach, live HEK293 cells were either stained with membrane and nuclear dyes or expressed Actin-GFP (Fig. 3J-L, Fig. S3E). Pre- and post-BaM imaging revealed fluorescence signal localization in both live cells and BaM particles (Fig. 3J-K, Fig. S3E). Line-scan intensity profiles of individual particles further confirmed signal levels above background (Fig. 3L), indicating capture of cellular material per nano-particle.

To assess the versatility of BaM, we applied this technique to live HEK293 cells expressing various GFP-tagged cytoskeleton-associated proteins previously known to form condensates or aggregates during over-expression. These included CLIP170 [44][45][43][46], Tau-3R [47][48], SALS[49][50], and ATIP3 domain 3 [51](false-colored; Fig. 3M-O). Protein size, construct species, and GFP position are detailed in Fig. S3G. Pre-BaM imaging revealed rapid condensate or aggregate formation in live cells (Fig. 3M), whereas post-BaM gels displayed GFP-positive particles corresponding to each protein tested (Fig. 3N). Analysis of BaM captured nanoparticles via imaging confirmed successful collection of protein nano-droplets sitting on individual nano-particles, often towards one edge of the BaM particle (Fig. 3O).

In parallel to experiments in HEK293 cells, we tested BaM on a live non-model organism lacking genetic tools. For this we chose *Chaos (Pelomyxa) carolinensis*, a single-celled giant amoeba measuring 1-5 mm [52][53][54] with well characterized sol-gel transitions in the cytoplasm[55]. BaM particles capture cytoplasm efficiently in this large amoeboid cell (Fig. S3F), demonstrating the method’s applicability across diverse cell types.

Across all conditions presented, single BaM shots recovered nuclear (Fig. S3E), membrane, cytosolic, and condensate material directly from live cells in a single penetration event. This demonstrates BaM’s versatility in sampling diverse biomolecules directly from living cells while preserving spatial context and compatibility with fluorescence-based confirmatory analyses.

## Structural studies: Cryo-EM reveals BaM particles with membrane-enclosed cellular material

Fluorescence microscopy showed only BaM particles carrying previously known and pre-tagged cytosolic and membrane content (Fig. 4A). Next, to further examine these membranous and total unlabeled material from live cells, we used TEM (Fig. S4) and Cryo-EM (Fig. 4) on single BaM particles, enabling high-resolution ultrastructural characterization of BaM-captured cytoplasmic and membrane compartments wetting the gold surface of capture particles.

**Figure 4:**
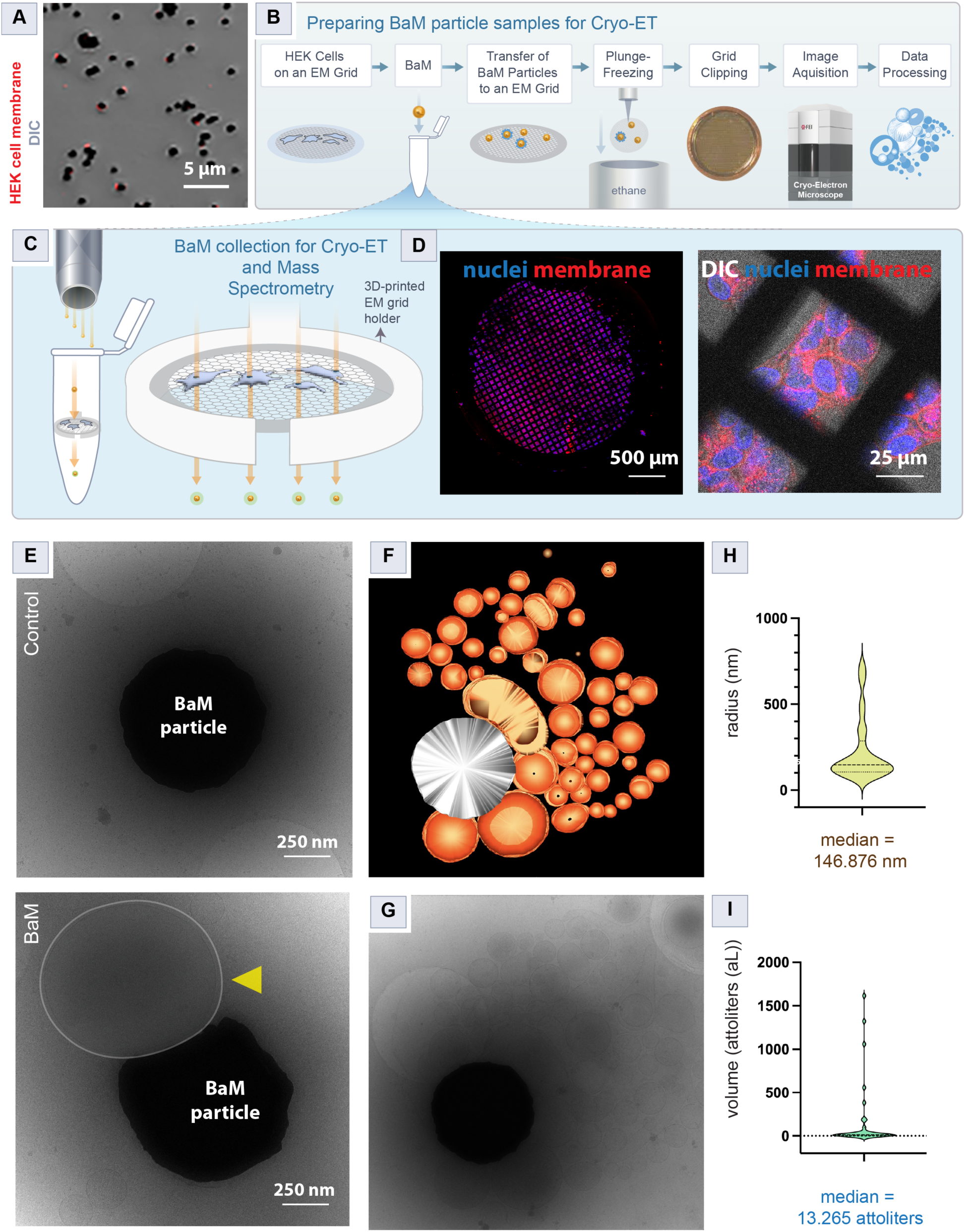
Structural analysis of BaM particles using Cryo-ET. A) Merged DIC and fluorescence image of BaM particles with membrane signals (red). (B) Schematic outlining the pipeline for preparing BaM particle samples for Cryo-ET. HEK293 cells are cultured on gold EM grids mounted in a custom 3D-printed holder inside a tube (C). After BaM, the particles are transferred to a Cryo-ET grid, plunge-frozen, clipped, imaged, and followed by image processing. (C) Collection setup with 3D-printed holder designed to house an EM grid inside a tube. (D) Fluorescence images of HEK293 cells cultured on an EM grid prior to BaM. Cells are labeled with membrane and nuclear markers. (E) Representative Cryo-EM micrograph of a single BaM particle. Top: control BaM particle without cellular material. Bottom: BaM particle with electron-dense membrane-enclosed cellular content (yellow arrowhead). (F) 3D reconstruction from a tilt series of a single BaM particle (white) using Cryo-ET. Individual volumes (orange spheres) represent segmented cellular cargo collected during the BaM shot (see Supplementary Video 3). (G) Unprocessed Cryo-EM micrograph used for 3D reconstruction. (H-I) Quantification of cellular material from micrographs. (H) Radius measurement and (I) estimated volume of collected material per BaM particle. Median values are indicated, calculated from 31-50 structures.

To enable direct high-resolution EM imaging of BaM particles, we immobilized HEK cells using a slightly modified method (Fig. 4B). The EM grid with live cells attached was transferred to a 3D-printed grid holder inside a collection tube (Fig. 4C), and cells were BaM imaged from above. This configuration enabled direct collection of penetrating BaM particles for subsequent electron microscopy (Fig. 4B). Because the time interval between BaM particle collection in the tube and transfer to the cryo-EM grid could allow cellular material to nonspecifically bind to BaM particles, we tested this possibility by collecting penetrating particles directly into liquid TEM resin in the collection tube (Fig. S4C). After resin solidification with embedded BaM particles, TEM imaging of sectioned resin confirmed the presence of cellular material attached to individual BaM particles (Fig. S4).

For Cryo-EM, BaM particles from the collection tube (see Methods) were transferred onto a quantifoil Cryo-EM grid and followed the Cryo-ET pipeline (Fig. 4B). Electron micrographs in Fig. 4E and Fig. S5 depict control BaM particles without cellular content alongside those containing captured material; arrows mark membrane-enclosed structures. In addition to 2D images, tilt-series of an individual BaM particle carrying cellular material were acquired and reconstructed in 3D (Fig. 4H-I; Supplementary Video 3). These electron micrographs revealed the architecture of the collected cellular content, which consisted primarily of spherical, membrane-bound structures with electron-dense lumens surrounded by single, double, or multi-lamellar membranes (Fig. S4; Fig. 4H-I). Cytosolic wetting was observed on BaM particles for some cytoplasmic droplets. Quantitative analysis indicated an average radius of ∼146 nm and a volume of ⇡13 attoliters per capture droplet (Fig. 4F-G). Given the ⇡4 pL volume of a typical cultured mammalian cell [56], this corresponds to ∼200,000 capture volumes of ∼20 attoliters each from a single cell. These results indicate that BaM can sample intact material from live cells including cytoplasmic and membrane components, making it well suited for high-resolution label-free ultrastructural studies of subcellular assemblies and protein complexes.

## BaM enables isolation and proteomic characterization of CLIP170 condensates

The capacity to sample subcellular material from live cells using BaM motivated us to investigate protein condensates formed by CLIP170 (Cytoplasmic Linker Protein of 170 kDa), a microtubule cytoskeleton-associated protein [57][58][59][60]. CLIP170 is well-characterized for its microtubule plus-end tip tracking [57][58] and for forming protein condensates in cells and in vitro through liquid-liquid phase separation (LLPS) [44][43]. Here, we revisited CLIP170 condensate and determined its protein composition by coupling BaM particles with mass spectrometry. Because the sensitivity of mass spectrometry currently is insufficient to detect cellular content from a single BaM particle (attoliter scale), we pooled GFP-CLIP170 samples collected from a large number of individual BaM particles (see Methods for details). Pooling was achieved using magnetic beads conjugated with anti-GFP antibodies, which selectively bind GFP-tagged proteins (Fig. 5A), only picking BaM particles with GFP-CLIP170.

**Figure 5:**
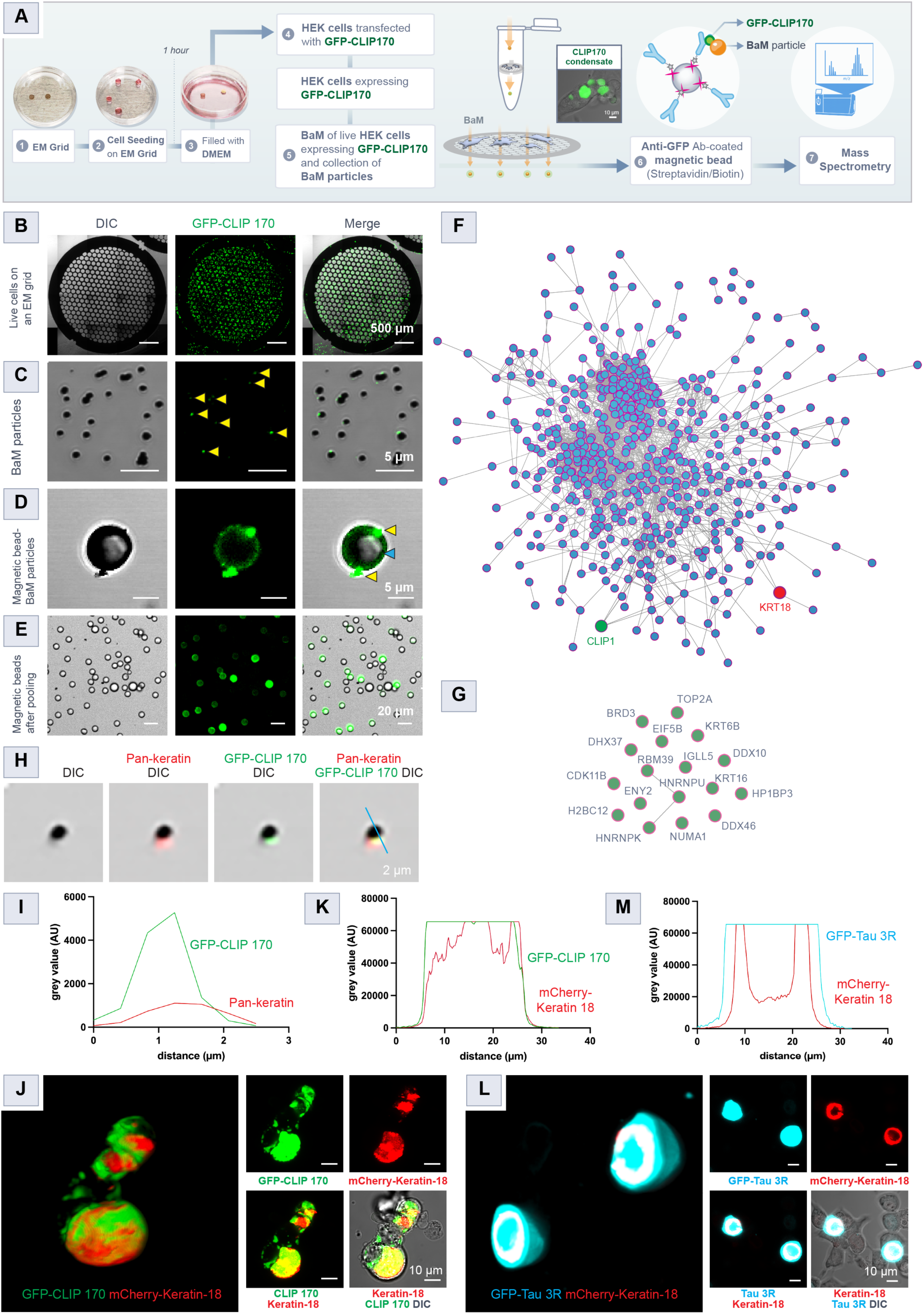
BaM enables the collection and proteomic analysis of CLIP170 condensates, revealing an association between Keratin and both CLIP170 and Tau condensates. (A) Schematic showing the pipeline for the collection of CLIP170 condensates and GFP (control) from live HEK293 cells using BaM. Cells cultured on EM grids expressing GFP-CLIP170 or GFP and subjected to BaM using the collection setup described in Fig. 4B-C. Collected BaM particles were pooled with anti-GFP-coated magnetic beads and analysed using mass spectrometry (see Methods). (B-E) Representative images from key steps in the pooling protocol. (B) GFP-CLIP170-expressing cells on an EM grid. (C) BaM particles showing captured GFP-CLIP170 signal (yellow arrowheads). (D) Magnetic bead (blue arrow) coated with anti-GFP antibody binds GFP-CLIP170-bound BaM particles (yellow arrow). (E) Image of isolated magnetic particles used for mass spectrometry. n = 2 independent experiments. (F) PPI network of 641 proteins identified in the CLIP170 condensate, generated from mass spectrometry data using the STRING database. Each blue node represents a protein, and connecting grey lines indicate previous experimentally validated interactions. CLIP1(CLIP170) and KRT18 are highlighted in green and red. See Fig. S7 for protein names with interaction networks, and Supplementary Table 2 for the full list of proteins. (G) GFP control BaM experiment: PPI network of 18 proteins identified from magnetic beads incubated with BaM particles penetrated through HEK293 cells expressing only GFP. (H) BaM particles showing colocalization of GFP-CLIP170 (green) with pan-Keratin antibody staining (red). n=3. (I) Line scan analysis confirms spatial overlap of the two signals. (J-K) Confocal images and corresponding line scans from live cells co-expressing GFP-CLIP170 and mCherry-Keratin-18 confirm their co-localization. n = 102 cells positive for CLIP170 and Keratin. See also Fig. S8 and Supplementary Video S4. (L-M) Co-expression of Tau 3R-GFP (cyan) and Keratin-18-mCherry (red) in live cells reveals a distinct cage-like organization of Tau 3R condensates around Keratin. (M) Line scan confirms this organization. n = 110 cells. See also Fig. S8 and Supplementary Video S5.

HEK293 cells expressing GFP-tagged CLIP170 proteins were BaM imaged as previously described (Fig. 4B, Fig. 5A-B). Fluorescence imaging post BaM confirmed successful collection of GFP-CLIP170 condensates (Fig. 5C) on individual BaM particles. To isolate only these condensates, freshly penetrated BaM particles were incubated with 8*µ*m magnetic beads coated with anti-GFP antibodies (Fig. 5D; Fig. S6A). After washing away non-specifically bound cell content, the magnetic beads were used for mass spectrometry. Although many BaM particles detached during washing, intact GFP-CLIP170 condensates remained on magnetic beads for downstream analysis (Fig. 5E; Fig. S6B). A protein gel of this BaM sample showed a band corresponding to GFP-CLIP170 (Fig. S6C).

Across multiple mass spectrometry replicates, we identified 641 proteins enriched in the pooled GFP-CLIP170 condensates (Supplementary Table 2; Fig. 5F, Fig. S7A-B). STRING-based protein-protein interaction (PPI) analysis [61] revealed a highly interconnected Protein-Protein Interaction (PPI) network (Fig. 5F; Fig. S7A-B), suggesting that CLIP-170 condensates are heterogeneous mixtures of proteins and their interacting partner proteins. Classification of the 641 proteins using the PANTHER (Protein ANnotation Through Evolutionary Relationship) system (www.pantherdb.org) [62] revealed a broad range of cellular functions (Fig. S6D).

As controls, BaM particles without penetrating cells yielded no detectable proteins, and BaM particles penetrating cells expressing only GFP identified 18 proteins, with 101 overlapping between the GFP control and the GFP-CLIP170 condensates (Fig. 5G).

Together, these results establish BaM as a new method for isolating intact protein condensates from live cells for proteomic analysis, enabling the discovery of previously unrecognized protein associations.

## Discovery of Keratin intermediate filaments association with CLIP170 and Tau condensates

Although, mass spectrometry identified 641 proteins associated with CLIP170 condensates, forming an extensive PPI network (Fig. 5F), we detected a correlation of Keratin-18 consistently detected in CLIP170 condensates across experiments (Fig. 5F).

Given the presence of Keratin-18 in the CLIP170 interactome, we next explored spatial relationship of this unique cytoskeleton component in both the BaM particle (Fig. 5H; Fig. S8E) and in live cells (Fig. 5J; Fig. S8G-I). Immunofluorescence staining of BaM particles revealed co-localization of GFP-CLIP170 with pan-Keratin antibody signals (Fig. 5H), confirmed by line-scan analyses showing overlapping intensity profiles (Fig. 5I-K; Fig. S8F). Live-cell confocal imaging of HEK293 cells co-expressing GFP-CLIP170 and mCherry-Keratin-18 demonstrated a unique co-localization of Keratin-18 within condensates (Fig. 5J-K; Fig. S8H-I; Supplementary Video S4). In contrast, co-expression of GFP-CLIP170 and mCherry-Keratin-14 showed orthogonal localization (Fig. S8I; Supplementary Video S4), establishing Keratin-18 indeed is preferentially assembles large structures inside CLIP170 condensate droplets.

To test whether Keratin association extended to other cytoskeletal condensates, we examined Tau, a microtubule-associated protein that stabilizes axonal microtubules, involved in axonal transport, and is implicated in neurodegenerative tauopathies such as Alzheimer’s disease [63][64][65][66][67].Tau undergoes condensation both in vitro and in vivo, highlighting its intrinsic propensity to form dense phases under physiological and experimental conditions [68][69][70]. In HEK293 cells co-expressing Tau 3R-GFP and mCherry-Keratin-18, confocal imaging revealed a striking cage-like onion-ring arrangement of Keratin-18 surrounding Tau 3R condensates (Fig. 5L). Line-scan analyses confirmed this spatial organization, with Keratin and Tau 3R signals forming complementary intensity profiles (Fig. 5M; Fig. S8J-K; Supplementary Video S5). This remarkable arrangement of a cytoskeletal component within a condensate body would have not been reveled without our unique label-free discovery approach enabled by BaM.

These results demonstrate that Keratin-18 is a consistent partner with both CLIP170 and Tau 3R condensates, suggesting a broader role for intermediate filaments in organizing and stabilizing condensate assemblies. Thus combined BaM-proteomics approach enables systematic mapping of condensate proteomes and their structural interfaces with cellular scaffolds.

## Discussion

We report Ballistic Microscopy (BaM) using nano-particles instead of photons as an imaging probe. This new method enables single-cell spatial biology and material isolation of unlabeled cytoplasmic components while preserving spatio-temporal arrangements leading to discovery and characterization of new subcellular structures in live cells. Traditional single-cell sampling methods often rely on bulk extraction, which disrupts spatial information [21][23][25]. In contrast, BaM provides spatially resolved, in situ retrieval of intracellular material from live cells, without the need for any genetic modification, fixation, or whole-cell lysis. The collected “physical image” embedded in a hydrogel “film” can then be analyzed by established downstream methods including anything from Mass Spectrometry or Cryo-electron Microscopy for structural and biochemical characterization to imaging for RNA-seq. This unique capability opens new avenues for studying anything from uncharacterized single molecules to multiprotein complexes, together with their interactomes, at single-cell or subcellular resolution in situ.

Using BaM, we successfully captured and analysed attoliter-scale cytosolic content (Fig. 4G), including protein condensates such as those formed by CLIP170 (Fig. 5A-G). Mass spectrometry of CLIP170 condensate identified 641 proteins (Table 1, Fig. 5F), which assembled into a dense protein-protein interaction network (Fig. S7A-B) encompassing a wide range of cellular functions (Fig. S7D-E). Strikingly, a large fraction of proteins associated with CLIP170 condensates were RNA-binding proteins, consistent with previous findings in condensate proteomics [71][72][73]. Furthermore, 17.73% (Fig. S7E) of total proteins present in CLIP170 condensates remain unclassified, representing the “dark proteome”[9] further highlighting the importance of label free method such as BaM. Decoupling capture and analysis into two steps alleviates constraints on experiments - opening the possibility of conducting destructive experiments on live cells without compromising temporal resolution.

Our unlabeled approach led to the discovery of Keratin 18 present in CLIP170 protein condensates. Such unmasking can enable traditional imaging techniques to provide further insights. In our case, Keratin 18 showed a distinct localization pattern with both GFP-CLIP170 and GFP-Tau 3R under the same live-cell conditions (Fig. 5J-L; Fig. 8H; Fig. 8J). In the case of Tau 3R, Keratin 18 formed a closed concentric shell-like structure around the nucleus, covered by Tau (Fig. S8J-K, Supplementary Video 5). While the role of cytoskeletal elements in condensate stabilization has been previously predicted [74][75], this represents the first experimental evidence of Keratin intermediate filaments co-localized within condensates. Functional characterization or ultrastructural analysis of Keratin filaments lies beyond the scope of this study but could be addressed in the future using Keratin-targeting drugs, knockouts, CRISPR approaches, or high-resolution imaging methods such as Cryo-EM or Cryo-CLEM.

In our fluorescence microscopy and structural analyses of single BaM particles containing cellular material, we consistently observed that captured contents were not evenly distributed but instead localized toward one edge of the particles (Fig. 3L, Fig. 3O, Fig. 4E; Fig. S4E; Fig. S5) - akin to adhesion of a fluid droplet on a high curvature surface of the particle. This asymmetric distribution is a classical example of surface wetting of protein droplets with a finite contact angle [76]. Cryo-ET further revealed that most captured cellular contents were membrane-enclosed structures with high electron-dense regions, often appearing as single or multiple membrane layers (Fig. S4E; Fig. S5). Further theoretical studies will allow us to understand the complete regime of membrane tube formation that enables these cytoplasmic droplets to escape the cell.

Despite the need for further optimization and refinement, the current BaM technique represents a platform with immense potential for a new label-free method for understanding spatial distribution of molecules within a living cell - both in native and diseased state. By enabling direct “structure-to-context” correlation, BaM bridges a critical gap between imaging-based localization and biochemical or structural analysis. Future efforts will focus on developing and improving the temporal resolution of BaM, allowing repeated and long term sampling of live cells to isolate and study dynamic structures over time. Future version of BaM can also include specific targeting of individual subcellular structures including cilia, spindles, filopodia, organelles, membranes, and disease-associated protein condensates or aggregates, while also capturing previously unknown proteins across different cell types. Beyond live-cell samples, BaM also holds the potential to extend to tissue scales or thick brain slices where photons or electrons cannot penetrate, including in non-model organisms. Importantly, systematic spatial mapping of protein interactomes with BaM could generate comprehensive atlases of cellular organization. Coupled with artificial intelligence (AI), such experimental datasets may ultimately enable predictive models linking protein interactions to cellular functions and phenotypic outcomes, offering new insights into cellular homeostasis and disease mechanisms that have remained inaccessible.

## Limitations

Reflecting the native state of the cytoplasm, the cellular content collected by BaM particles is highly heterogeneous. By using small ballistic particles, we ensure that the collected cytoplasmic droplets are small in size ( 146 nm radius) and volume ( 13 attoliters). This is advantageous for single-cell spatial biology but currently also poses challenges for high-resolution molecular analysis with small volume of isolated material. To address the limited sample volume, enrichment strategies targeting specific molecules such as pooled sampling were implemented (Fig. 5A). While we demonstrate proteomic identification and protein-protein interaction network mapping of isolated CLIP170 condensates from live HEK293 cells (Fig. 5F), ultra-low abundance components may remain below detection thresholds of current techniques. Coupling BaM with more sensitive detection platforms, such as ultrasensitive MS workflows for attomole samples [77][78][79] or single-molecule protein and nucleotide sequencing approaches [80][81] will expand the detectable biomolecular range. Established methods for cell lysates [82][83] or single-cell bulk cytoplasm [84], could be adapted and customized for BaM.

Since samples are collected throughout the length of the traversal path, the current BaM system lacks Z-axis depth resolution. Thus BaM is currently suited for thin samples. A penetrating ballistic particle can collect material all along its trajectory inside a cell depending on the physical properties and surface chemistry of the gold particle. Direct visualization of membrane-labelled trails of individual BaM particles (Fig. 3I; Supplementary Video 2), together with previous theoretical studies [85], suggests the possibility of capturing molecules from different cellular depths along these tracks. However, this remains speculative, and careful experiments are required to extract this information.

The current BaM setup utilizes EM grids or Vitrogel hydrogel films which limit HEK cell adhesion. This creates a low yield for multiple time resolved BaM shots (low temporal BaM). Improving this adhesion efficiency and reducing the shock wave from the gas will further improve our capacity to do long term continous BaM imaging. Utilizing an alternative strategy for accelerating nano-particles, such as laser-induced projectile impact testing (LIPIT) [36][37][86], can generate high-speed particles without shockwave effects will enable long term BaM imaging. Advances in nanoparticle research offer potential solutions, with commercially available gold particles of varied composition, geometry, surface chemistry, or antibody conjugation that could mitigate stress effects [87][88]. Taken together, we provide a roadmap for the continued development of BaM. We believe BaM presents a new and versatile approach to sampling and identifying unknown bimolecular components and assemblies in living cells - shedding light on the dark proteome [89] of biology.

## Materials and Methods

### Custom Ballistic Microscopy setup

A custom-made BaM setup was constructed from scratch, consisting of: 1) a particle acceleration unit, 2) particle capture unit (hydrogel films) and 3) an integrated multi-channel imaging unit. The particle acceleration unit, inspired by previously reported open gene gun designs [27][28], operates on compressed helium gas (Fig. 2a, Fig. S1A-B). A high-pressure helium gas cylinder (HE UHP300, Airgas) was connected to an electronically controlled solenoid valve (H2B19-00Y) controlled with a time delay relay (Dayton 1EGB4, 50 ms) via a pressure gauge rated up to 1500 psi. The solenoid valve was attached to a detachable nozzle designed specifically to reduce shock waves during particle acceleration. The nozzle featured a double convergent-divergent tube construction with an open end and multiple pressure relief valves to minimize shock wave arriving to the sample of interest. The solenoid valve’s opening and closing operation was controlled with an 8-pin time-delay relay and a firing switch (Fig. 2b, Fig. S1A-B).

To position BaM gold particles above the sample, the particle acceleration unit was mounted vertically above the hydrogel film. Just before the pressure relief valves, a BaM particle-loading mechanism was integrated, consisting of an O-ring spacer, and a circular piece of parafilm (acting as a rupture film) to securely hold the BaM particles (Fig. 2a). All joints between components were carefully sealed with high-pressure-grade pipe fittings with Teflon tape (6802K22, McMaster) to ensure safety.

In addition to the particle acceleration unit, the BaM setup included a traditional imaging unit (Fig. 2a, Fig. S1A-B) consisting of a custom-built inverted fluorescence and brightfield microscope controlled by bespoke interface software [41], paired with a commercial high-speed camera (Phantom VEO 640S, 4788 fps). The entire BaM setup, except for the gas cylinder, was mounted inside a closed cabinet for safety and control. The cabinet was assembled on a vibration-free optical table to maintain stability during experiments (Fig. S1A-B). Detailed methods can be found in the supplementary materials.

## Acknowledgments

The authors acknowledge use of shared facilties at Stanford campus including Stanford Cell Sciences Imaging Facility (CSIF), the Vincent Coates Foundation Mass Spectrometry (RRID:SCR-017801) Thermo Orbitrap Eclipse nanoLC/MS system (RRID:SCR-022212), funded by NIH Shared facility (1S10OD030473), Stanford Cancer Institute Proteomics/Mass Spectrometry Shared Resource (NIH P30 CA124435 ), Stanford-SLAC CryoET facility (NIH U24 GM139166), Midwest Center for Cryo-ET (MCCET) and the Cryo-EM Research Center (NIH grant U24 GM139168). We thank Lydia-Marie Joubert for cryo-EM training and S. Garvey McKenzie for mass spectrometry training and advise. We sincerely thank Rebecca Konte for creatingschematic diagrams and providing artistic input for the figures. Authors thank current members of the Prakash laboratory for helpful discussions.

## Author contributions

A.S.J. and M.P. designed the research. A.S.J. performed experiments. R.C. and M.P. developed theoretical framework. R.C. developed simulations and scaling and 3D cryo-ET reconstruction video. A.S.J analyzed the data with discussion from M.P. A.S.J and R.C wrote the first draft. All authors revised and edited the paper.

## Competing interests

The BaM technology described in this work is covered by a published US patent (WO/2024/044276) assigned to M.P. and A.S.J. All authors declare no competing interests.

## Data and materials availability

All commercial resources and antibodies used here are detailed in Supplementary Table.

Supplementary materials

## Materials and Methods

Supplementary Text

Figures S1 to S8

Movie S1-S5

Table S1-S2

Captions for Movies S1 to S5

## Notes

### Competing Interest Statement

The authors have declared no competing interest.

