## Supplementary materials for "Ballistic Microscopy (BaM)"

### Supplementary Materials for
### Ballistic Microscopy (BaM)

A. S. Jijumon, Ray Chang, Manu Prakash†

†

**This PDF file includes:**

Materials and Methods
Supplementary Text
Figures S1 to S8
Table S1 to S2
Captions for Movies S1 to S5

**Other Supplementary Materials for this manuscript:**

Movies S1 to S5 (see links)

### Materials and Methods

#### Preparation of gold particles for BaM experiments

Solid gold micro- and nanoparticles with diameters of 600 nm, 1 $\mu$m, 1.6 $\mu$m, 2.5 $\mu$m, and 5.1 $\mu$m were utilized in this study. Although smaller ballaistic particles were also used, systematic data is presented for the above listed dimensions. For ease of detection and larger capture profile - 600 nm and 1 $\mu$m gold particles were primarily used for most of the protein capture assays. The particle sizes were verified using Transmission Electron Microscopy (data not shown). To minimize aggregation during storage, gold particles were prepared using a standardized protocol as detailed below. Gold particles of the desired diameters (600 nm, 1 $\mu$m, 1.6 $\mu$m, 2.5 $\mu$m, and 0.15 $\mu$m) were purchased (Bio-Rad Laboratories, USA) in powdered form. For each preparation, 10-20 mg of gold particles were weighed and transferred to a 1.5 mL microcentrifuge tube. The particles were suspended in 500 $\mu$L of 100% ethanol and vortexed for 2 minutes. The suspension was then centrifuged at 9391 g for 2 minutes. The supernatant was discarded, ensuring that a small volume of ethanol remained to keep the particles immersed. This washing and dispersion process was repeated three times. The prepared particles were stored in 100% ethanol at either 4°C or room temperature for up to one week before use in the BaM experiments.

#### Ballistic microscopy: preparation and operation

A custom particle-loading mechanism was developed to load particles on a rupture disk. A circular, clean piece of parafilm (diameter 1 cm) was cut and attached beneath an O-ring (inner diameter 6 mm), ensuring the gold nanoparticles deposited on the parafilm face inwards towards the helium gas line. The parafilm was isotropically stretched to ensure even rupture during firing. To maintain consistency in the number of BaM particles across experiments, 50-100 $\mu$g of gold particles in powder form was weighed and placed on the inward-facing side of the parafilm under the O-ring spacer. This assembly was then inserted into the hose fitting connected to the divergent-convergent nozzle. The nozzle was attached back to the particle acceleration unit, and the desired pressure was set using the regulator knob on the gas cylinder, with pressure ranging from 50 to 600 psi. The sample was positioned under the BaM nozzle tip, and the region of interest was aligned with the nozzle hole using 405 nm laser light from the inverted microscope. The distance between the nozzle tip and the sample was maintained at 40 mm. Once alignment was complete, particles were fired onto the sample by opening the solenoid valve using the firing switch. If required, the side view of particle penetration through the gel was captured using a modular high-speed camera (Phantom VEO 640S, 4000 fps), or the bottom view of the BaM process was imaged using the inverted fluorescence microscope (Squid multi-color epi-fluroescence Fig. 2A; Fig. S1A-B). For high-resolution imaging of the BaM particles, the samples were immediately transferred to a Zeiss LSM 780 inverted multiphoton laser scanning confocal microscope following BaM.

**BaM particle penetration depth experiments**

To measure the penetration depth of BaM particles (Fig. S2A-C), particles of varying sizes (600 nm, 1 $\mu$m, 1.6 $\mu$m, 2.5 $\mu$m, and 5.1 $\mu$m) were shot onto agarose gels of differing concentrations (0.5%, 1%, and 2%). For gel preparation, 0.5, 1, or 2 grams of low-melting agarose powder (Cat# BP165-25; Fisher scientific) was weighed and added to 100 mL of warm Milli-Q water in a conical flask. The mixture was heated in 20-40 second intervals in a microwave, swirling carefully between intervals, until the agarose was completely dissolved, forming a clear solution with no visible particles. Once dissolved, the solution was cooled to 50-60°C by leaving it at room temperature for 10-20 minutes. After cooling, 4 mL of the gel mixture was poured into 35 mm diameter petri dishes (Falcon, Cat# 351008), ensuring no bubble formation. The gel was then left to solidify at room temperature for 1-3 hours. Fully solidified gels were stored at 4°C and could be used for BaM experiments for up to one week.

BaM experiments were conducted using all combinations of particle sizes (600 nm, 1 $\mu$m, 1.6 $\mu$m, 2.5 $\mu$m, and 5.1 $\mu$m), pressures (50, 100, 150, and 200 psi) , and agarose gel concentrations (0.5%, 1%, and 2%) (Fig. S2A-C). After particles were shot into the gel, the gel was carefully sliced into thin sections (1-2 mm thick, parallel to the particle trajectory) using a clean single-edge razor blade (Electron Microscopy Sciences, Cat# 71962). The slices were then placed on a 22×40 mm, #1.5 coverslip (Fisherbrand, Cat# 12-544-B) for imaging (Fig. 2D; Fig. 2G-H; Fig. S1e). DIC imaging was performed using an inverted Nikon ECLIPSE Ti2 microscope equipped with a 4X DIC objective (Fig. S2A-C). The particle penetration depth was quantified using Fiji software by manually marking the particle positions relative to the gel surface. Graphs were generated using GraphPad-Prism software (Version 10.3.0) to analyze and visualize the results (Fig. S2D).

#### High-speed imaging of BaM particles

High-speed imaging of ballistic particles in gel was performed using a Phantom V1210 high-speed camera, capable of capturing images at speeds up to 64,000 fps. The camera was coupled with a 10x Mitutoyo objective and mounted perpendicular to the BaM particle trajectory axis, as shown in Fig. S1 B and F. A custom setup with an additional light source (Lambda XL, Sutter Instruments) was used to enhance contrast (Fig. 2E; Fig. S1G). Camera settings and data acquisition were controlled using Phantom software (Phantom Camera Control (PCC 3.6)).

For imaging 5.1 $\mu$m diameter particle penetration into the gel, a 1% agarose gel was prepared as described above and poured into the lid of an Ibidi dish (Ibidi, Cat#80826), creating a thin gel layer. The camera was focused on the edge of the dish, capturing the field of view at the air-gel interface (Fig. 2E; Fig. S1G; Supplementary video 1).

#### HEK 293 cell culture

HEK 293 cells (Human Embryonic Kidney cells; ATCC CRL-1573) were cultured in DMEM medium supplemented with 10% fetal bovine serum (FBS; Sigma-Aldrich), 2 mM L-glutamine (Life Technologies), and 1X penicillin-streptomycin (Pen Strep; Life Technologies). Cells were maintained in T25 flasks (Nunc™ EasYFlask™ Cell Culture Flasks) at 37°C with 5% $CO_2$ in a humidified cell culture incubator. Cultures were split when they reached 90% confluency, and the DMEM medium was replaced at least twice per week. For BaM experiments, cells were seeded and prepared either in a 3D hydrogel or on an EM grid, depending on experimental requirements described below.

#### Cell culture on EM grids for BaM

Gold (Electron Microscopy Sciences; Cat# CF-222C-10-AU) or molybdenum (PELCO; Cat # 5GM200) grids were used to attach HEK 293 cells for BaM experiments. A gold grid was carefully placed on a glass-bottom cell culture dish (WillCo Wells, Cat# GWST-5030) under a biosafety cabinet. Subsequently, 100 $\mu$L of HEK 293 cells ($10^6$ cells/mL) in DMEM medium was added to the EM grid as shown in the Fig. 5a. The medium containing cells was dispensed as a drop from the pipette tip onto the grid without allowing contact between the pipette and the grid. The cell density was adjusted based on the experimental requirements; lower cell density was used for spatially distinguishing BaM particles, while higher cell density was applied for Cryo-EM or mass spectrometry experiments requiring maximum BaM particle with cellular content. The cell culture dish containing the EM grid was transferred to a cell culture incubator and incubated for 1-2 hours to allow cell attachment. After this initial incubation, the dish was gently filled with DMEM medium and incubated for an additional 12-24 hours. Cell health and density were checked using a dissection microscope before proceeding with BaM experiments. If required, live cell staining for specific markers, such as nuclei, cell membranes, or cytoskeleton components, was performed directly on the cells attached to the EM grid. Transfection with proteins of interest could be conducted either on the grid or by seeding pre-transfected cells onto the grid.

##### Cell culture on hydrogel for BaM

VitroGel HEK293 (The Well Bioscience, Cat# VHM05) was used to attach HEK 293 cells for BaM experiments. The hydrogel was prepared following the manufacturer's guidelines, with modifications specific to BaM experiments. VitroGel, stored at 4°C, was first brought to room temperature or warmed at 37°C for 10 minutes. It was then mixed with DMEM medium in a 2:1 (v/v) ratio. A glass-bottom 24-well plate was used for the BaM setup, with 500 $\mu$L of the gel mixture added to each well. The plate was gently swirled to ensure an even coating of gel on the bottom of the well, as uneven coverage could allow cells to grow beneath the gel by escaping through gaps between the gel and the well wall. The gel was incubated in a cell culture incubator at 37°C for 20 minutes. After the initial incubation, 500 $\mu$L of DMEM medium was gently added along the walls of the well to avoid disturbing the gel. The plate was incubated for 12 hours in a cell culture incubator to allow complete gel polymerization. Following this, 500 $\mu$L of cell culture medium containing HEK 293 cells ($5\times10^5$ cells/mL) was added to the well. The cells were allowed to grow for 12-24 hours, and their health and attachment were checked using a dissection microscope before proceeding with BaM experiments. Cells were transfected with plasmid encoding proteins of interest before adding to the hydrogel. However, live cell staining for markers such as membranes, nuclei, or cytoskeleton was performed while the cells were on the gel. Extended incubation of HEK 293 cells on hydrogel promotes the formation of cell spheroids. Under these conditions, the BaM parameters must be adjusted to achieve greater particle penetration depths. Optionally, HEK 293 cells could be mixed directly with the VitroGel-DMEM solution during the gel formation process, embedding the cells within the hydrogel as it solidified. However, adding cells on top of the hydrogel provided sufficient depth for BaM particles to penetrate effectively.

##### Cell transfection and live staining for BaM

Actin-GFP, GFP-CLIP170, GFP-Tau 3R, SALS-GFP, and GFP-ATIP3 domain 3 constructs [43] were used to transfect HEK 293 cells to validate the BaM approach for collecting cellular materials. Details of the construct are shown in Fig. S3G. HEK 293 cells at approximately 80% confluency in DMEM medium were transfected using Lipofectamine 3000 Transfection Reagent (Invitrogen, Cat# L3000001), following the manufacturer's guidelines. GFP expression levels were checked 24 hours post-transfection using a fluorescence microscope. For proteins involved in phase separation or aggregate formation, BaM was conducted 48-72 hours post-transfection to ensure adequate protein expression.

For cells attached to EM grids, transfection was performed directly on the grid-bound cells. In contrast, for cells grown on VitroGel hydrogel or polycarbonate membranes, pre-transfected cells were seeded to maximize transfection efficiency.

For live-cell staining, HEK 293 cells were stained with 0.1 $\mu$g/mL of Hoechst 33342 (Invitrogen, Cat# H3570) to visualize nuclei and CellMask plasma membrane stain ( Invitroge, Cat# C10045; 1:500) for cell membranes. The stains were diluted in DMEM medium and incubated with the cells for 20 minutes. After staining, the medium was replaced with fresh cell culture medium, and the staining was verified using a fluorescence microscope before conducting BaM experiments (Fig. 3, 4).

##### *Chaos (Pelomyxa) carolinensis*

*Chaos (Pelomyxa) carolinensis* culture was obtained from Carolina biological supply (Cat# 7646630), and maintained according to the provided instructions. The culture jar was stored at room temperature with the lid loosely placed over the jar's mouth to allow air exchange. No additional feeding was required prior to BaM experiments. Before performing BaM, the organisms were stained with the live-cell marker CellTrace ( Thermo Fisher Scientific, Cat# C34564) for 20 minutes, following the manufacturer's guidelines. Their activity and mobility were confirmed using a fluorescence microscope to ensure they were healthy and active after staining. For BaM, the organisms were placed on a thin layer of 1% agarose gel on a glass-bottom dish with minimal liquid to immobilize them. BaM was conducted using 1 $\mu$m gold particles, and the gel was immediately imaged using a Zeiss LSM 780 inverted multiphoton laser scanning confocal microscope with a PLAN APO 63x oil objective (NA 1.4).

**BaM experiments using a GFP-actin gel layer and DNA amplification on BaM particles**

Concentrated and clarified cell lysate containing GFP-actin was prepared from HEK-293 cells using a previously published method [43][83]. A 1% agarose gel was prepared in a 35 mm diameter petri dish as described earlier. Following this, 40 $\mu$L of GFP-actin lysate was carefully pipetted onto a ~5 mm diameter area on the gel. The lysate-covered area of the gel was aligned with the BaM nozzle, and BaM particles with a diameter of 5.1 $\mu$m were fired onto the lysate (Fig. 2F-H). After BaM, the gel was carefully cut into thin slices (Fig. S1E) and imaged the GFP fluorescence using a Zeiss LSM 780 microscope as described above (Fig. 2F-H; Fig. S2E-F). A control experiment was performed using the same protocol, except the BaM particles were fired onto agarose gel without any cell lysate (Fig. S1F).

To test DNA collection on BaM particles from the surface to the interior of the gel, and to test direct DNA amplification on the particles, experiments were performed using a 10% acrylamide gel (Fig. S2H). For this, a total of 5 mL of gel solution was prepared by mixing 1.65 mL of acrylamide/bis-acrylamide (29:1), 1.25 mL of 1M Tris–HCl (pH 8), 20 $\mu$L of TEMED, 20 $\mu$L of 10% ammonium persulfate (APS), and 2.06 mL of Milli-Q water. Subsequently, 4 mL of this mixture was poured into a 35 mm diameter glass-bottom dish and allowed to polymerize at room temperature for 2 hours. The master mix for DNA amplification on BaM particles was composed of the following components: 50 $\mu$L of WarmStart LAMP 2X Master Mix (ID: E1700, New England Biolabs), 2 $\mu$L of 1 U/$\mu$L Uracil-DNA Glycosylase (ID: EN0361, Thermo Fisher), 0.25 $\mu$L of 100 mM dUTP solution (ID: R0133, Thermo Fisher), 2 $\mu$L of LAMP Fluorescent Dye (ID: B1700S, New England Biolabs), 4 $\mu$L of 3 mM HNB dye (Vienna BioCenter), 10 $\mu$L of a primer mix (specific to *Schistosoma mansoni* as a test case), 20 $\mu$L template (5ng/ $\mu$L) and 31.75 $\mu$L of molecular biology-grade water, resulting in a total reaction volume of 120 $\mu$L. A 50 $\mu$L volume of the prepared master mix was pipetted onto the solidified acrylamide gel within an approximate 5 mm diameter area. The targeted region on the gel was aligned with the BaM nozzle and bombarded with 5.1 $\mu$m diameter BaM particles. After particle penetration, the gel was immediately transferred to a 65 °C dry bath and incubated for 60 minutes [90]. DNA amplification on the BaM particles was evaluated by observing the fluorescence signal on the particles embedded within the gel using a Zeiss LSM 780 confocal fluorescence microscope (Fig. S2H).

#### Ballistic Microscopy of HEK293 cells cultured on EM grids and gels

*Ballistic Microscopy of EM grids with cells on sodium acrylate gel:*

To preserve the spatial information of penetrated BaM particles, EM grids with live HEK293 cells were placed on a sodium acrylate-based gel prior to BaM (gel composition: [91]) (Fig. 3A-I). The unpolymerized gel solution was prepared using 23% (w/v) sodium acrylate, 10% (w/v) acrylamide, 0.1% (w/v) N,N′-methylenebisacrylamide (bis-acrylamide), and 1× PBS. This solution can be aliquoted and stored at -20°C. Before BaM, the gel was prepared as follows: 2.5 $\mu$L of 10% tetramethylethylenediamine (TEMED) and 2.5 $\mu$L of 10% ammonium persulfate (APS) were added to 45 $\mu$L of the thawed unpolymerized gel solution on ice. Then, 20 $\mu$L of the gel solution was pipetted onto a clean glass-bottom dish and covered gently with a 12 mm round coverslip using forceps. Care was taken to ensure an even distribution of the gel across the radius. The assembly was incubated at 37 °C for 1 hour to allow polymerization. After polymerization, the coverslip was carefully removed with forceps, leaving a thin, transparent gel layer on the glass-bottom dish. The gel must be freshly prepared immediately before BaM. To minimize gel shrinkage over time, a piece of wet Kimwipe (Cat #06-666) can be placed inside the dish without touching the gel.

To perform BaM on cells cultured on EM grids, the BaM setup was first prepared by loading the gold particles and setting the appropriate pressure. The EM grid containing HEK 293 cells (cultured as described above) was carefully removed from the petri dish using tweezers, and excess DMEM liquid was gently removed by touching the grid edge to a clean Kimwipe. The EM grid with cells was then placed on the polymerized gel for BaM (Fig. 3A-B; Fig. S3A). Following the BaM, the grid was carefully removed from the gel to proceed with downstream processing (Fig. 3c). High-magnification imaging of embedded BaM particles in the gel was performed either using the built-in fluorescence microscopy unit of the BaM setup or transferred to a commercial confocal microscope (Fig. 3).

*Ballistic Microscopy of EM grids with cells for mass spectrometry and Cryo-EM:*

The EM grid with live cells was inserted into a custom 3D-printed EM grid holder designed to fit inside a 200 $\mu$L PCR tube (Fig. 4B-C; Fig.5A). The PCR tube containing the grid holder was placed into a bread board stage base (Thorlabs, Cat# 973/579-7227) pre-chilled at -20 °C for 1-3 hours. For sample pooling, the same grid holder could be reused for experiments involving the same cell type and particle size. For temperature-sensitive protein samples, the tube was immediately transferred to ice before proceeding with the subsequent steps, depending on experimental requirements.

*Ballistic microscopy of cells cultured on VitroGel HEK293 hydrogel:*

Before performing BaM on cells cultured on VitroGel hydrogel (Fig. 3J-N), the BaM setup was prepared for firing as previously described. The petri dish containing the hydrogel with attached cells was removed from the cell culture incubator, and the DMEM medium was carefully aspirated to remove as much as possible. The dish was then aligned under the nozzle tip of the BaM setup using the laser from the inverted microscope. A fixed distance of 40 mm was maintained between the nozzle tip and the hydrogel surface to ensure consistency. The cells were immediately bombarded with BaM particles by activating the firing switch. Following the experiment, fresh DMEM medium was added to the petri dish, and the microparticles carrying cellular content were imaged using a Zeiss LSM 780 inverted multiphoton laser scanning confocal microscope (Fig. 3J-N).

#### Transmission electron microscopy of BaM particles

$10^6$ HEK 293 cells were seeded and grown on gold EM grids (Electron Microscopy Sciences; Cat# CF-222C-10-AU) for 12-24 hours as described above (Fig. S4A-C). After preparing the BaM setup for firing, the EM grid was placed into a custom EM grid holder inside a 200 $\mu$L PCR tube filled with Epon resin (EMS, Hatfield, USA) (Fig. S4C). The resin volume was adjusted to ensure that the grid holder did not come into contact with the liquid resin. The cells were bombarded with 0.6-1 $\mu$m diameter gold particles at 100 psi pressure, as described above. After firing, the grid holder containing the grid was carefully removed using tweezers, and the PCR tube containing the resin and BaM particles were transferred to a 65 °C oven and incubated for 16-20 hours to allow the resin to solidify [92]. Once the resin was solidified, the block containing the BaM particles with cellular content was sectioned using a Leica UC6 ultramicrotome. The PCR tube was carefully removed, and the sides of the resin block were trimmed using a single-edge razor blade (Electron

Microscopy Sciences, Cat# 71962). The sample was mounted, and 1 $\mu$m-thick sections were cut using a diamond knife (Ultra 45°; DiATOME diamond knives). These thicker sections ensured that 1 $\mu$m BaM particles were included within the slices (Fig. S4C). The sections were collected onto formvar-coated 100-mesh copper grids ( Electron Microscopy Sciences, Cat# FCF100H-Cu). The sections on the grids were stained with 2% uranyl acetate and Reynold's lead citrate for 5 minutes each at room temperature. Subsequently, the stained sections were imaged using a JEOL JEM-1400 transmission electron microscope operating at 120 kV, and images were captured with a Gatan Orius 2k × 2k digital camera (Fig. S4D-E). All image processing and analysis were performed using Fiji software.

**Cryo-electron microscopy of BaM particles**

8 $\mu$L of DMEM medium containing 106 HEK 293 cells/ mL were seeded and grown on one Molybdenum EM grid as described above. After 12-24 hours the cells were bammed using 1 $\mu$m diameter gold particles in a 200 $\mu$L PCR tube, 10 $\mu$L of cold PBS supplemented with 1X Protease inhibitor cocktail (PIC; GenDepot, Cat# P3100-001) was added to the particles. A Quantifoil gold holey carbon grid R 1.2/1.3 were glow-discharged in a GloQube Glow Discharge (Quorum Technologies) system for 90 s. Sample application and vitrification were carried out with a Leica Em GP2 Plunge Freezer set to 25 °C and 100% humidity, and blotted for 3 seconds using a Whatman No. 1 filter paper.

Cryo-EM data were collected on a Titan Krios G4i transmission electron microscope (ThermoFisher Scientific) operated at 300 kV and equipped with a C-FEG (cold field emission gun), a SelectrisX energy filter (10 eV slit width), and a Falcon 4i direct electron detector (ThermoFisher Scientific). Automated data acquisition was performed using SerialEM v4.0. Data were acquired at a nominal magnification of 64,000×, corresponding to a calibrated pixel size of 1.965 Å per pixel at the specimen level. A total electron dose of 80 electrons/$\text{Å}^2$ was distributed over the tilt series, which was acquired using a dose-symmetric tilt-scheme covering a range from +60° to -60° in 3° increments with a group size of 2. The nominal defocus range was set between -3 $\mu$m and -5 $\mu$m. A 100 $\mu$m objective aperture and a 70 $\mu$m C2 aperture were used during imaging.

*Cryo-electron tomography reconstruction*

Tilt series were aligned and reconstructed using the Etomo graphical interface from the IMOD software package (v5.1.1; University of Colorado Boulder). Fiducial-less alignment was performed using cross-correlation-based patch tracking across all tilt images. No binning was applied during alignment or reconstruction, and the full-resolution images (1.965 Å/pixel) were used throughout. Weighted back-projection (WBP) was employed to compute the 3D tomographic volume, with the reconstructed tomogram spanning 4096 × 4096 × 500 voxels. The tilt axis angle was automatically estimated by Etomo from image metadata. After reconstruction, structures of interest were segmented manually in 3dmod using interpolated contours, with visual inspection across Z-slices to confirm continuity.

#### Mass spectrometry of CLIP170 condensates isolated from live cells using BaM

*Ballistic Microscopy and pooling of CLIP170 condensates:*

Due to the lower protein sensitivity of the mass spectrometer used, BaM was performed on 75-100 EM grids per experiment, which were subsequently pooled to ensure adequate cellular material for analysis. For this, 8 $\mu$L of DMEM medium containing $10^6$ cells/mL was seeded onto each EM grid as described above (Fig. 5A). After 12-24 hours from cell seeding, HEK 293 cells were transfected with GFP-CLIP170 plasmids using Lipofectamine 3000 transfection reagent following the manufacturer's instructions. Transfected cells were incubated for 24-48 hours to allow for protein expression. Before performing BaM, the transfected cells on EM grids were checked under a fluorescence microscope to confirm strong GFP fluorescence (Fig. 5B). EM grid batches with poor transfection efficiency or weak GFP signals were discarded and the experiment restarted with new batch of grids to avoid low yield during immunomagnetic isolation and subsequent mass spectrometry.

Before starting BaM experiments, streptavidin magnetic beads (Bangs Laboratories, Cat# UMC0102) were conjugated with a biotinylated anti-GFP antibody (Abcam, Cat# ab6658)[93]. To prepare the conjugated beads, 40 $\mu$L of magnetic beads were washed three times with 10X volume of cold 1X PBS. For each wash, the beads were resuspended by gentle tapping and pelleted by centrifugation at 9391$g$ for 3 minutes. After the final PBS wash, the beads were resuspended to maintain a 40 $\mu$L slurry volume. Subsequently, 12 $\mu$L of biotinylated anti-GFP antibody was added, and the mixture was incubated for 1-2 hours at room temperature with gentle shaking to allow for conjugation. After conjugation, unbound antibodies were removed by washing the magnetic beads three times with 250 $\mu$L of cold 1X PBS, pelleting the beads by centrifugation at 9391$g$ for 3 minutes for each wash. Following the final wash, the beads were resuspended to a final volume of 40 $\mu$L. The beads were kept on ice until the next step.

Following preparation of the conjugated magnetic beads, BaM was performed on EM grids containing HEK 293 cells expressing GFP-CLIP170 condensates. All BaM particles were pooled into a single 200 $\mu$L PCR tube setup, as showed in Fig. 5A. First, the culture dish containing the EM grid was taken out of the cell culture incubator. Using tweezers, the grid was gently blotted with a clean filter paper to remove residual DMEM medium before inserting into a 3D-printed grid holder inside the PCR tube. The cells were bombarded with 1 $\mu$m gold particles as described previously. This process was repeated for all grids. Before the BaM experiments, the PCR tube was chilled using a cold a bread board stage base (Thorlabs, Cat# 973/579-7227), which was pre-cooled at -20°C for 2-3 hours. During grid exchanges and BaM setup preparation, the tube was consistently kept on ice to minimize potential protein degradations. After collecting the BaM particles, 100 $\mu$L of cold PBS supplemented with 1X protease inhibitor cocktail was added to the PCR tube. The BaM particles were then transferred to a new tube containing the conjugated magnetic beads. The mixture was incubated for 6-10 hours at 5 rpm at 4°C to allow binding of GFP-tagged proteins to the antibody-conjugated beads. Before and after adding the magnetic beads, 1 $\mu$L of BaM particles was examined under a fluorescence microscope for initial validation (Fig. 5C; Fig. S6A). Following incubation, the magnetic beads were separated using a magnetic rack (250 $\mu$L tubes; Sergi Lab Supplies). The supernatant was discarded and replaced with 150 $\mu$L of fresh cold PBS containing 1X PIC. This washing step was repeated 5-10 times to reduce nonspecific protein binding to the magnetic beads. The washed magnetic beads were either directly used for LC-MS/MS analysis (Fig.5A,E) or prepared for SDS-PAGE followed by Coomassie Brilliant Blue staining (Fig. S6C). For control pull-down and mass spectrometry experiments, the same procedure was followed using HEK 293 cells transfected with a GFP plasmid instead of GFP-CLIP170 (Fig. 5G).

*CLIP170 Sample Preparation:*

BaM protein samples on magnetic beads (CLIP170) or gold nanoparticles (control) were digested. The proteins were reduced by 500 mM of dithriothreitol (DTT), added to a final concentration of 10mM and incubated for five minutes at 55°C, followed by head-over-head mixing at room temperature for 25 minutes using a Barnstead Thermolyne Labquake rotisserie shaker. They were then allowed to cool and alkylated with 1M acrylamide, added to a final concentration of 30 mM, at room temperature for 30 minutes. This was followed by overnight digestion at 37°C using 500 ng of mass spectrometry grade trypsin/LysC mix (Promega). Post-digestion, samples were quenched with formic acid (adjusted to a pH ~3) and desalted using MonoSpin C18 Solid-Phase Extraction (SPE) columns (GL Sciences). Finally, the samples were dried via SpeedVac (ThermoFisher Scientific) and exchanged into LC-MS reconstitution buffer (2% acetonitrile with 0.1% formic acid in water) for instrumental analysis.

*LC-MS/MS Analysis:*

Proteolytically digested peptides were separated using an in-house pulled and packed reversed phase analytical column (~25 cm in length, 100 microns of I.D.), with Dr. Maisch 1.9 micron C18 beads as the stationary phase. Separation was performed with a 95-minute reverse-phase gradient (2-35% B, followed by a high-B wash) on an Acquity M-Class UPLC system (Waters Corporation) at a flow rate of 450 nL/ min. Mobile Phase A was 0.2% formic acid in water, while Mobile Phase B was 0.2% formic acid in acetonitrile. Ions were formed by electrospray ionization and analyzed by an Orbitrap Eclipse Tribrid mass spectrometer (Thermo Scientific). The mass spectrometer operated in a data-dependent mode, using CID fragmentation.

*Data Analysis:*

The RAW data were analyzed using Byonic v5.5.2 (Protein Metrics) to identify peptides and infer proteins. A concatenated FASTA file containing Uniprot *Homo sapiens* proteins, bait sequences, and other likely contaminants and impurities was used to generate an *in silico* peptide library. Proteolysis with Trypsin/LysC was assumed to be fully specific. The precursor ion tolerance was set to 12 ppm. The fragment ion tolerance was set to 0.4 Da. Cysteine modified with propionamide was set as a fixed modification in the search. Variable modifications included oxidation on methionine, dioxidation on tryptophan, deamidation of glutamine and asparagine, and cyclization of glutamine and glutamic acid. Proteins were held to a false discovery rate of 1% using the standard reverse-decoy technique [94].

#### Immunostaining of BaM particles post-BaM and fluorescence imaging

After BaM, the hydrogel containing embedded BaM particles was first blocked with 1% Bovine Serum Albumin (BSA) (Fisher Scientific, Cat# BP9703-100) prepared in 1× PBS for 20 minutes at room temperature. The blocking solution was then replaced with the primary antibody solution, prepared in 1% BSA blocking buffer supplemented with 0.1% TritonX-100 (Sigma-Aldrich, Cat# 9036-19-5). For immunostaining, anti-GFP antibody (Thermo Fisher Scientific; Cat# A-11122; 1:100 in 1X PBS) (Fig. S8B), and YL1/2 anti-tyrosinated tubulin antibody (Sigma-Aldrich; Cat# MAB1864-I; 1:100 in 1X PBS) (Fig. S8D) were incubated for 30 minutes, while Pan-Keratin antibody (Thermo Fisher Scientific; Cat# 41-9003-82; 1:100in 1X PBS) was incubated for 4 hours (Fig. 5H; Fig. S8E-G). Following incubation, the samples were washed three times with 1X PBS supplemented with 0.1% Triton X-100, with each wash lasting 3 minutes and performed with gentle shaking. The secondary antibodies, Alexa 565-labeled anti-rat antibody (Thermo Fisher Scientific, Cat#A-11077) for YL1/2, and anti-rabbit Alexa 568 (Thermo Fisher Scientific, Cat# A-11011) for anti-GFP, were prepared in 0.1% BSA in 1X PBS (1:100). The samples were incubated with the antibodies for 20 minutes, followed by three washes with PBST (1X PBS with Triton X-100), as previously described. The stained BaM particles were imaged using a Zeiss LSM 780 inverted multiphoton laser scanning confocal microscope equipped with a PLAN APO 63X oil objective (NA 1.4) (Fig. 5H, Fig. S8E-G).

HEK293 cells expressing GFP-CLIP170 were immunostained with a pan-Keratin antibody (Fig. S8G) using an adapted version of a published protocol [95]. Cells were cultured on glass-bottom dishes (Pelco, Ted Pella Inc.) and transfected to express GFP-CLIP170 for 36 hours. After removing the culture medium, cells were gently rinsed with 1× PBS and fixed with 4% paraformaldehyde (Electron Microscopy Sciences) in 1× PBS for 20 minutes at room temperature. After fixation and PBS washes, cells were permeabilized (0.5% Triton X-100, 5 min) and incubated overnight at 4 °C with eFluor 570-conjugated pan-Keratin antibody (1:200) in 1% BSA. The samples were washed three times with PBST, mounted in Vectashield Antifade Mounting Medium (Vector Laboratories), and imaged using confocal microscopy.

##### Image and data analysis

Microscopy images were analyzed using Fiji or Zeiss ZEN Black software, and processed figures were assembled and annotated using Adobe Photoshop and Illustrator. Quantitative data were subsequently transferred to GraphPad Prism software for plotting and statistical analyses. Line scans of fluorescent signals on BaM particles were obtained by manually drawing lines across the particles (Fig. 2I; Fig. 3L,O; Fig. 5I,K,M; Fig. S2G,I; Fig. S8F). All data were then transferred to GraphPad Prism for plotting and statistical analysis. The penetration depth of BaM particles from the gel surface under different conditions (Fig. S2D) was measured by manually drawing lines for each particle using ImageJ, and the resulting graphs were also generated using GraphPad Prism. All supplementary videos were prepared using Adobe After Effects.

Protein-protein association networks (Fig. 5F,G, FIG S7a,b) derived from mass spectrometry data were analyzed using custom VBA macros in Microsoft Excel and the STRING database (https://string-db.org/). Classification of the 641 proteins identified in the GFP-CLIP170 condensate was performed using the PANTHER (protein annotation through evolutionary relationship) classification system (www.pantherdb.org) (Fig. S6D).

### Supplementary Text

#### Theoretical considerations

The design of ballistic microscopy is not only of engineering interest but also of theoretical significance. Very few studies have examined the physical processes of biolistics, as most studies have focused on optimizing the delivery of genetic materials[96, 97]. A study by Zhang et al. proposed the first physical analysis for the biolistics process[98]. However, they primarily addressed the aerodynamics of the particles before tissue penetration, and modeled the particle-tissue interactions in an over simplistic manner. Another study by Zhang et al. used molecular dynamics simulations (MD-sim) to delineate the physical interactions between high-speed particles and biological membranes[31]. Nonetheless, their simulation did not include hydrodynamics interactions, as the particles travel in a vacuum. An intuitive approach that incorporates the hydrodynamic interactions between high-speed nanoparticles and living cells is still lacking. Here, we use scaling analysis guided by past MD-sim results to provide key physical insights into the feasibility of this novel cytoplasm sampling method.

**Physical regime of Ballistic Microscopy** The high-speed nanoparticles in BaM exist in a unique regime, defined by small size, extreme velocity, and interfacial interactions. Depending on the particle delivery methods, particle velocities can reach up to 3000 m/s[99], which falls into the supersonic regime even in water. Based on the typical parameter values associated with the process (length scale $L$ ~ particle diameter $D$ ~ $1\mu$m, fluid density $\rho_\mathrm{f}$ ~ 1000 kg/m$^3$, interfacial tension $\sigma$ ~ 50 mN/m[29], velocity scale $U_0$ ~ 1000 m/s, dynamic viscosity $\mu$ ~ 1 mPa-s, speed of sound $c$ ~ 1500 m/s, gravitational acceleration $g$ ~ 9.8 m/s$^2$), we find that BaM is a rare example of a low-Bond-number, high-Weber-number, high-Reynolds-number, high-capillary-number, and intermediate-to-high-Mach-number interfacial phenomenon (Table S1).

Compared to other high-speed penetrating interfacial phenomena (Fig. 1C), BaM falls into a unique region of the Weber-Bond number phase space. While most other penetrating phenomena lie along the diagonal, BaM is distinctly off-diagonal in a low-Bond-number, high-Weber-number quadrant. We can understand this intuitively by looking at the ratio between Weber number and Bond number. As the interfacial tension term cancel out with each other, the ratio of the two becomes $\mathrm{We}/\mathrm{Bo} = U_0^2/(gL) \sim (aL)/(gL) \sim a/g$, where $a$ is the characteristic acceleration assuming the particle is accelerated for a length comparable to its own length. The fact that microparticles in BaM is off-diagonal from other penetrating interfacial phenomena indicates its extraordinarily high acceleration in its mechanism.

These dimensionless numbers also suggest the dominant physical forces during particle deceleration. In most interfacial phenomena, deceleration arises from buoyancy, interfacial tension, viscous forces, or inertia. For microscale systems like phage peneration or microinjection, viscous and interfacial forces dominate. In macroscale cases like bullets hitting water, inertia and buoyancy dominate. BaM uniquely lies in a regime where only inertial forces are significant in the deceleration process.

**Table S1**: Physical regime of Ballistic Microscopy.

| dimensionless number | Bond number | Weber number | Reynolds number | Capillary number | Mach number |
|---|---|---|---|---|---|
| symbols | Bo | We | Re | Ca | Ma |
| physical meaning | $\frac{\text{gravity}}{\text{interfacial tension}}$ | $\frac{\text{inertia}}{\text{interfacial tension}}$ | $\frac{\text{inertia}}{\text{viscosity}}$ | $\frac{\text{viscosity}}{\text{interfacial tension}}$ | $\frac{\text{speed}}{\text{speed of sound}}$ |
| definition | $\rho_\mathrm{f} g L^2/\sigma$ | $\rho_\mathrm{f} U_0^2 L/\sigma$ | $\rho_\mathrm{f} U_0 L/\mu$ | $\mu U_0/\sigma$ | $U_0/c$ |
| values | $2 \times 10^{-7}$ | $2 \times 10^4$ | $10^3$ | 20 | 0.67 |

Another unique aspect of BaM is its extreme separation in both time and length scales (Figs. 1D&E). These gaps make a complete theoretical analysis extremely challenging, requiring multiscale approaches. For example, the difference between membrane healing time (~ 10 min) and membrane-crossing time for a 1 $\mu$m at 1000 m/s (~ 5 ps) spans 14 orders of magnitude. Similarly, inertial deceleration and membrane crossing occur on time scales much shorter than membrane bending or protein binding. This indicates that forces from membrane bending and specific surface chemistry (e.g., antibody or streptavidin coatings) play less role during cytoplasmic sampling. Spatially, the lipid bilayer thickness (~4-5 nm) is several orders of magnitude smaller than a typical mammalian cell (~10-20$\mu$m). This massive length scale separation also creates a simulation challenge, as resolving all relevant scales simultaneously would render computational modeling prohibitively expensive.

**Physical intuitions about the deceleration process** Despite these challenges, some physical intuitions can still help us understand the feasibility of BaM. To achieve cytoplasm sampling, a particle must fully traverse the cytoplasm and reach the gel beneath the cell. If the particle cannot reach the bottom membrane, sampling fails. Thus, assuming the cytoplasm has water-like viscosity gives an upper bound for the penetration length. The particles decelerate via fluid drag, described by:

$$\begin{aligned} \frac{\mathrm{d}v}{\mathrm{d}t} &= -\frac{3}{8} c_{\mathrm{D}}(\mathrm{Re}) \frac{\rho_{\mathrm{f}}}{\rho_{\mathrm{p}}} \frac{v^2}{R} \\ \Delta x(t) &= \int_0^t v(t')\mathrm{d}t' \end{aligned} \tag{S1}$$

where $v$ is the particle velocity, $c_{\mathrm{D}}$ is the drag coefficient using Brown & Lawler's correlation[30], $\rho_{\mathrm{p}}$ is the particle mass density, $R$ is the particle radius, and $\Delta x$ is the traveling distance of the particle. Using Equation S1, we estimate the penetration depth and compare it with typical cell height (~3-4 $\mu$m). As shown in Fig. 1F, particles $\geq$ 200 nm can easily reach the cell base with velocity of 200 m/s. Particles of 100 nm, on the other hand, require speeds $\geq$ 1000 m/s to penetrate the cell. As the spatial resolution of our imaging technique is ultimately limited by projectile size, this analysis indicates that the theoretical spatial resolution limit is comparable to typical light microscopy techniques.

As the friction between high-speed particles and water intrinsically generates heat, our approach could raise the concern of heating. The particle's peak temperature results from a balance between frictional heating and heat loss into the surrounding cytoplasm, described by coupled drag-interfacial thermal conductance equations:

$$\begin{aligned}\frac{\mathrm{d}v}{\mathrm{d}t} &= -\frac{3}{8}c_{\mathrm{D}}(\mathrm{Re})\frac{\rho_{\mathrm{f}}}{\rho_{\mathrm{p}}}\frac{v^2}{R},\\ \frac{\mathrm{d}(\Delta T)}{\mathrm{d}t} &= -\frac{v}{c_{\mathrm{p}}}\frac{\mathrm{d}v}{\mathrm{d}t} - \frac{3(\Delta T)(\mathrm{ITC})}{Rc_{\mathrm{p}}\rho_{\mathrm{p}}},\end{aligned} \tag{S2}$$

where $c_{\mathrm{p}}$ is the specific heat capacity of the particle, $\Delta T$ is the temperature raise, and ITC is the interfacial thermal conductance (~200 MW/m$^2$-K for water-gold interface[100]). For a 1 $\mu$m particle traveling at 1000 m/s, the peak temperature elevation is about 200K. However, as the water-gold interface has a relatively good thermal conductance, the temperature elevation rapidly decays to near baseline within 0.5 $\mu$s (Fig. 1G). Additionally, the peak temperature rise decreases with particle size because particle heat capacity scales with $R^3$ and interfacial area scales with $R^2$, both contributing to thermal moderation (Fig. 1H).

BaM's extreme velocities also raise concerns about cavitation, which can be evaluated by the cavitation number:

$$\mathrm{Cav} = \frac{p_{\mathrm{a}} - p_{\mathrm{v}}}{{}^{1}\!/_{2}\rho_{\mathrm{f}}v^2}, \tag{S3}$$

where $p_{\mathrm{a}}$ is the ambient pressure and $p_{\mathrm{v}}$ is the water vapor pressure. Cavitation is expected when $\mathrm{Cav} < 0.1$, and supercavitation when $\mathrm{Cav} \ll 0.1$.[101] At BaM speeds, cavitation and even supercavitation are likely (Fig. 1I). However, even though cavitation may occur, it does not mean the cavitation bubble could grow large because the rapid traveling time of the particle also limits the time when surrounding fluid stays under low pressure. Balancing the two effects, we can estimate the maximum radius of the cavitation bubble to be $R_{\mathrm{cav,max}} \approx 2D(-\mathrm{Cav} - C_{\mathrm{pmin}})$. For a 1 $\mu$m size particle, the maximum radius of its cavitation bubble is only 0.2 $\mu$m (Fig. 1J).

**Physical intuitions about particle-membrane interactions** In the previous section, we discussed the physical regime and important phenomena associated with the deceleration process of a BaM particle. However, as BaM intrinsically involves membrane perforation, in this section we discuss the importance of the particle-membrane interactions and what we know based on past simulation results.

As mentioned above, Zhang et al. used molecular dynamics simulation (MD-sim) to delineate the physical interactions between high-speed particles and biological membranes[31]. They used a solvent-free membrane model and bombarded it with nanoparticles. In their simulation, they identified three physical regimes. At low speed, the particle cannot penetrate the membrane and is rebounded by the interfacial tension of the membrane. At intermediate speed, the membrane forms a tube and is eventually penetrated by the particle, with the membrane attached. At high speed, the particle directly penetrates the membrane, with limited membrane content tagging along, and the membrane subsequently seals itself.

While their simulation results and identified physical regimes are insightful, they did not account for any hydrodynamic interactions, as the particles travel in a vacuum in their simulations. This omission makes it difficult to interpret the results in the context of BaM, where the particle–membrane interactions on the bottom membrane are more directly related to the amount of cytoplasm that can be extracted. Furthermore, BaM particles are surrounded by cytoplasm and experience significant drag forces.

Therefore, in addition to the three regimes proposed by Zhang et al. (rebound, tube formation, and direct penetration), we consider additional regimes that can only arise when the drag forces from the surrounding fluid are taken into account.

We first consider a regime that we call the self-stopping regime. In this regime, the particle cannot travel a distance equal to its own size because of fluid drag, and the particle does not require the particle-membrane interaction to stop. This regime can be easily identified by solving the Stokes drag equation and setting the maximum translation distance equal to the particle radius. The resulting critical velocity for the self-stopping regime of a particle traveling in a fluid is $v_{\text{crit}} = 9\mu/(2R\rho_{\text{p}})$. This means that any particle traveling at a speed lower than this critical velocity will stop even without interacting with the membrane.

We next consider the rebound regime of the membrane. To determine the velocity at which the particle will rebound off the membrane, we first calculate how fast a particle must move to penetrate the membrane, since all particles that fail to penetrate must rebound. Most biological membranes can only tolerate an area expansion of $\varepsilon$ ~3-5% before rupturing[102]. However, only the membrane in the vicinity of the particle penetration site will be involved, as the traveling time across the membrane is much shorter than the membrane stretching relaxation timescale or membrane diffusion timescale (Fig. 1D). Assuming the membrane area involved in the stretching process during particle penetration is about $\pi(D + 2\delta)^2$, where $\delta$ is the boundary layer thickness (see Section "Physical intuitions about the sampling process") around the particle, the particle must have enough kinetic energy to stretch the cell membrane beyond rupture. This can be written as the inequality:

$$\frac{1}{2}K_{\mathrm{A}}(\Delta A)^2 = \frac{1}{2}K_{\mathrm{A}}(\pi(D + 2\delta)^2\varepsilon)^2 < \frac{1}{12}\pi D^3\rho_{\mathrm{p}}v^2, \tag{S4}$$

where $K_{\mathrm{A}}$ is the area expansion modulus of a membrane, assumed to be 0.2 N/m[29]. From this inequality, we can calculate the critical velocity for membrane rupture as a function of particle diameter. The results recapitulate the simulations from Zhang et al.

In the simulation of Zhang et al., they found that when the particle velocity exceeds ~40 mm/s, the particle enters the direct penetration regime. This critical velocity is independent of the particle size. Based on the simulation parameters, we found that this critical velocity of 40 mm/s is comparable to the thermal velocity of the membrane (~ 11 mm/s). This suggests that when the thermal velocity of lipids is not fast enough to keep up with the particle velocity, the particle will directly penetrate the membrane locally. If the velocity of the particle falls between the critical velocity for particle rebound and the critical velocity for direct penetration, the membrane will undergo tube formation.

As mentioned earlier, BaM particles are immersed in fluid when they attempt to penetrate the bottom membrane of the cell. Therefore, the particle must still fall into the appropriate velocity regime after it has traveled a distance equal to its own size. Once we account for this fluid drag effect, we can construct a new phase diagram for particle–membrane interactions, based on Zhang et al.'s results and incorporating hydrodynamic effects (Fig. 1K). In the updated phase diagram, we find that the tube formation regime collapses into a very narrow region and is technically unachievable from an engineering perspective. The unknown regime in the diagram stems from the ambiguity in the thermal velocity of phospholipid molecules within the membrane. Therefore, most successful cytoplasm sampling would require the particle–membrane interaction at the bottom surface to fall within the direct penetration regime.

**Physical intuitions about the sampling outcome** From the above discussions, it is evident that successful sampling generally requires the bombarding particle to operate in the direct penetration regime. Consequently, the volume of cytoplasm that can be extracted is dominated by the volume of fluid that experiences momentum transfer from the high-speed particle.

The thickness of the fluid layers that undergoes momentum transfer can be characterized by the boundary layer thickness $\delta$. Considering the time over which the fluid experiences acceleration from the particle as $\tau = D/v$, the boundary layer thickness around the particle can be approximated as $\delta \sim \sqrt{2\nu\tau} = \sqrt{2\nu\frac{D}{v}}$, where $\nu$ is the kinematic viscosity of the cytoplasm. Based on this, we can obtain a naive estimation of the volume extracted as $V_{\text{ext}} \sim \pi D^2\delta \propto D^{5/2}\nu^{1/2}v^{-1/2}$. This shows that the extracted volume is highly dependent on the diameter of the particle. Thus, increasing the spatial resolution of BaM (i.e., using smaller particles) would require a corresponding improvement in the detection sensitivity of extracted biomolecules.

Using this naive scaling law for estimating extracted cytoplasmic volume, we can assess the feasibility of BaM for detecting various biomolecules. For a 1 $\mu$m particle penetrating the bottom membrane at a speed of 500 m/s, and assuming the cytoplasm has a kinematic viscosity of $10^{-6}$ $m^2$/s (a typical value for aqueous solution), the boundary layer thickness is about 63 nm. This yields an extracted volume of roughly 0.2 $\mu m^3$. Based on the typical protein content per mammalian cell (250 pg/cell)[32], averaged protein molecular weights (50 kDa)[32], the typical volume of a mammalian cell (4000 $\mu m^3$)[33], and the typical numbers of different kinds of protein in an organism ($10^4$ kinds of protein)[34], we estimate that in a 0.2 $\mu m^3$ volume, there are approximately $2 \times 10^5$ protein molecules and roughly 20 copies of a specific protein. As for the mRNA transcripts, based on the typical number per mammalian cell ($10^5 - 10^6$ mRNA/cell)[35], we estimate that a 0.2 $\mu m^3$ volume contains approximately 0.5-5 mRNA molecules. This indicates that BaM has the potential to achieve single mRNA transcript resolution.

**Summary** In summary, we have provided a comprehensive and intuitive scaling analysis that addresses several important physical considerations for BaM and its feasibility as a cytoplasm sampling method. We show that BaM can be characterized as a submicron-scale, high-Reynolds-number, high-Weber-number, high-Capillary-number, and low-Bond-number interfacial phenomenon. Its physics is dominated by fluid inertia and is marked by large separations in space and time scales. The entire ballistic process occurs in under 1–2 $\mu$s and involves transient heating and cavitation. Assuming a naïve scaling law based on boundary-layer theory and direct membrane penetration, a typical 1 $\mu$m BaM particle can extract cytoplasmic content containing proteins and mRNA within the detection limits of current state-of-the-art techniques.

### Supplementary figures

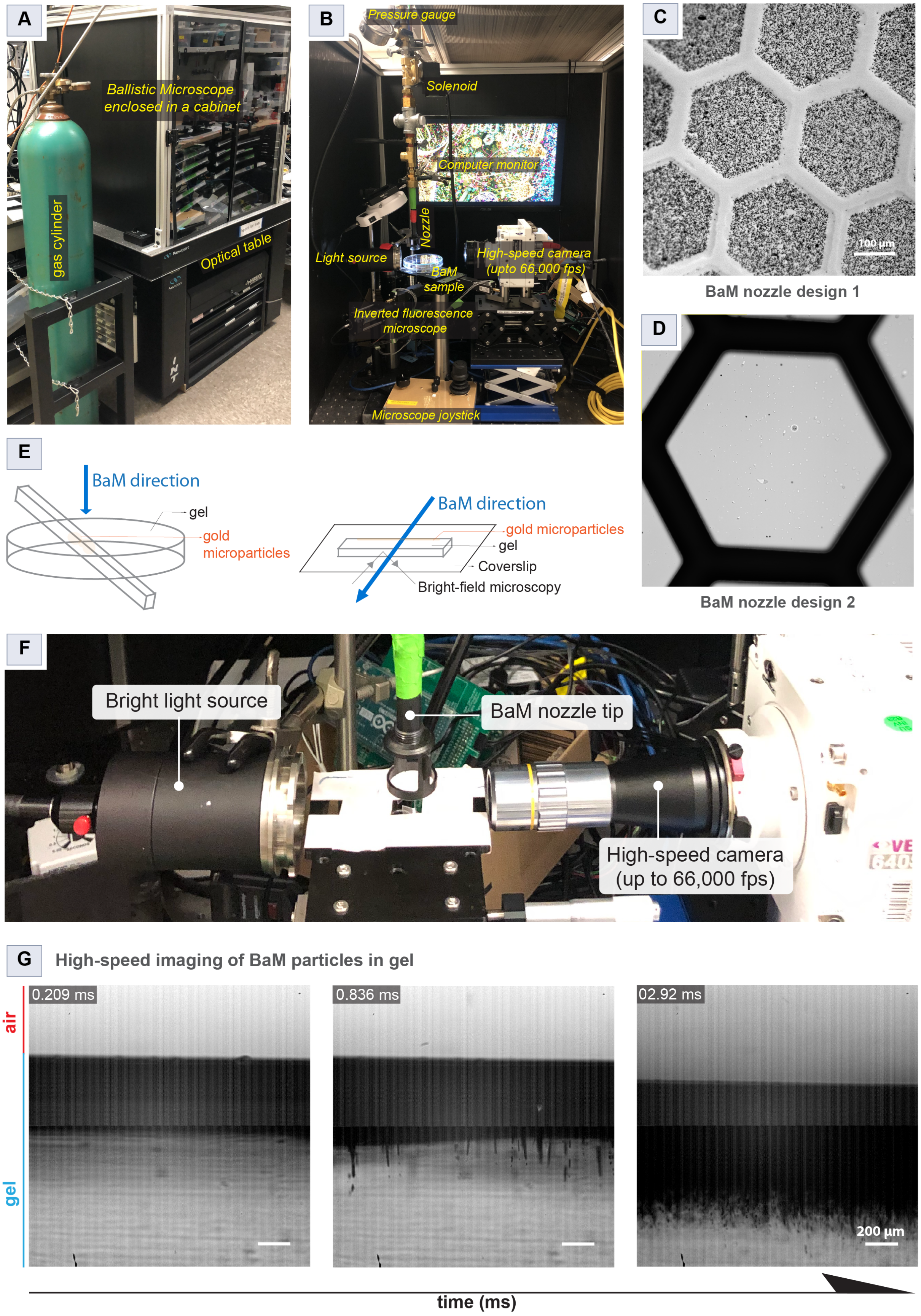




**Figure S1**: (Caption next page.)

**Figure S1**: **BaM instrument design, setup, and particle penetration characterization (complementary to Fig. 2).** (A-B) Photographs of the BaM setup enclosed in a custom cabinet on an optical table. Key components include a compressed gas cylinder, solenoid valve, pressure gauge, BaM nozzle, high-speed camera (up to 66,000 fps), inverted fluorescence microscope, and a computer. (C-D) Comparison of two BaM nozzle designs for particle delivery with reduced cell damage from the shockwave generated by the air puff. Sample destruction from the shockwave was tested by bombarding BaM particles into a hexagonally holed EM grid placed on an acrylate gel. The EM grid used here has a thin carbon layer. Bombarding with nozzle design 1 (C) leads to complete destruction of the layer due to the shockwave and dispersion of particles across the grid, which acts as a stencil. In contrast, design 2 (D) enables BaM particle penetration within the hexagonal holes without rupturing the carbon layer (magenta arrowheads). (E) Schematic of the sectioning and imaging method used to obtain a cross-sectional view of the gel for quantifying the penetration depth of gold microparticles. After bombardment, the gel was carefully cut using a clean blade. The cut section was then gently placed on a coverslip with the BaM penetration axis parallel to the coverslip surface (see also Methods). (F) Side view of the high-speed imaging configuration showing alignment of the BaM nozzle and camera with illumination source. (G) High-speed bright-field image sequence showing gold particle trajectories in 1% agarose gel post-BaM (20 psi), with timestamps in milliseconds. The interface between air (red) and gel (blue) is indicated. See Supplementary Video 1.

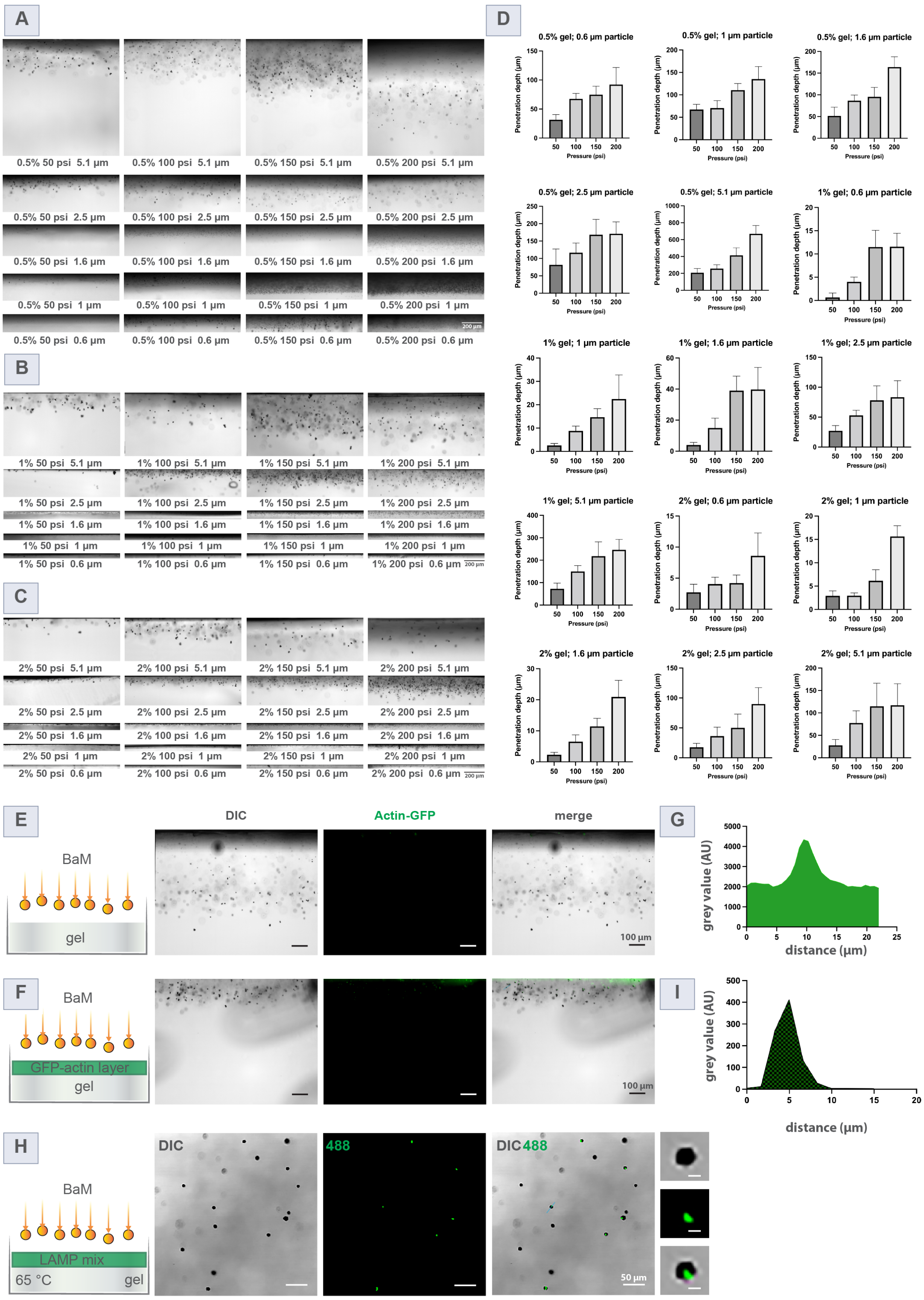




**Figure S2**: (Caption next page.)

**Figure S2**: **Quantification of BaM particle penetration and demonstration of DNA amplification on BaM particles.** (A-C) DIC images of gel cross-sections, prepared as shown in (Fig.S1e), under varying conditions: gel concentrations (0.5%, 1%, and 2%), particle diameters (0.6 $\mu$m, 1 $\mu$m, 1.5 $\mu$m, 2.5 $\mu$m and 5.1 $\mu$m), and pressures (50, 100, 150, 200 psi). (D) Quantification of penetration depth as a function of pressure and particle size across three gel concentrations. Data shown as mean ± SD. (E-G) Control for BaM-mediated protein pickup experiments (corresponding to Fig. 2F-I). In the absence of a lysate layer (E), no actin-GFP signal is detected on the penetrated particles. When a clarified HEK cell lysate expressing actin-GFP is layered above the gel (F), actin-GFP signals are observed on BaM particles, indicating successful protein pickup (cropped region of Fig. 2G-H). (G) Line scan analysis of a BaM particle (blue line in F) confirms the presence of a GFP signal. (H-I) A proof-of-concept experiment demonstrating the collection and amplification of DNA on 5.1 $\mu$m BaM particles using loop-mediated isothermal amplification (LAMP). A 1% agarose gel was coated with LAMP reagents and bombarded from the top with 5.1 $\mu$m microparticles, as shown in the schematic diagram (H) Subsequent incubation of the gel at 65 °C (see Methods), followed by fluorescence imaging, revealed DNA amplification on the BaM particles (bottom view of the gel). (I) Line intensity profile of fluorescence on an individual BaM particle (blue line in H) showing distinct peak corresponding to successful DNA collection.

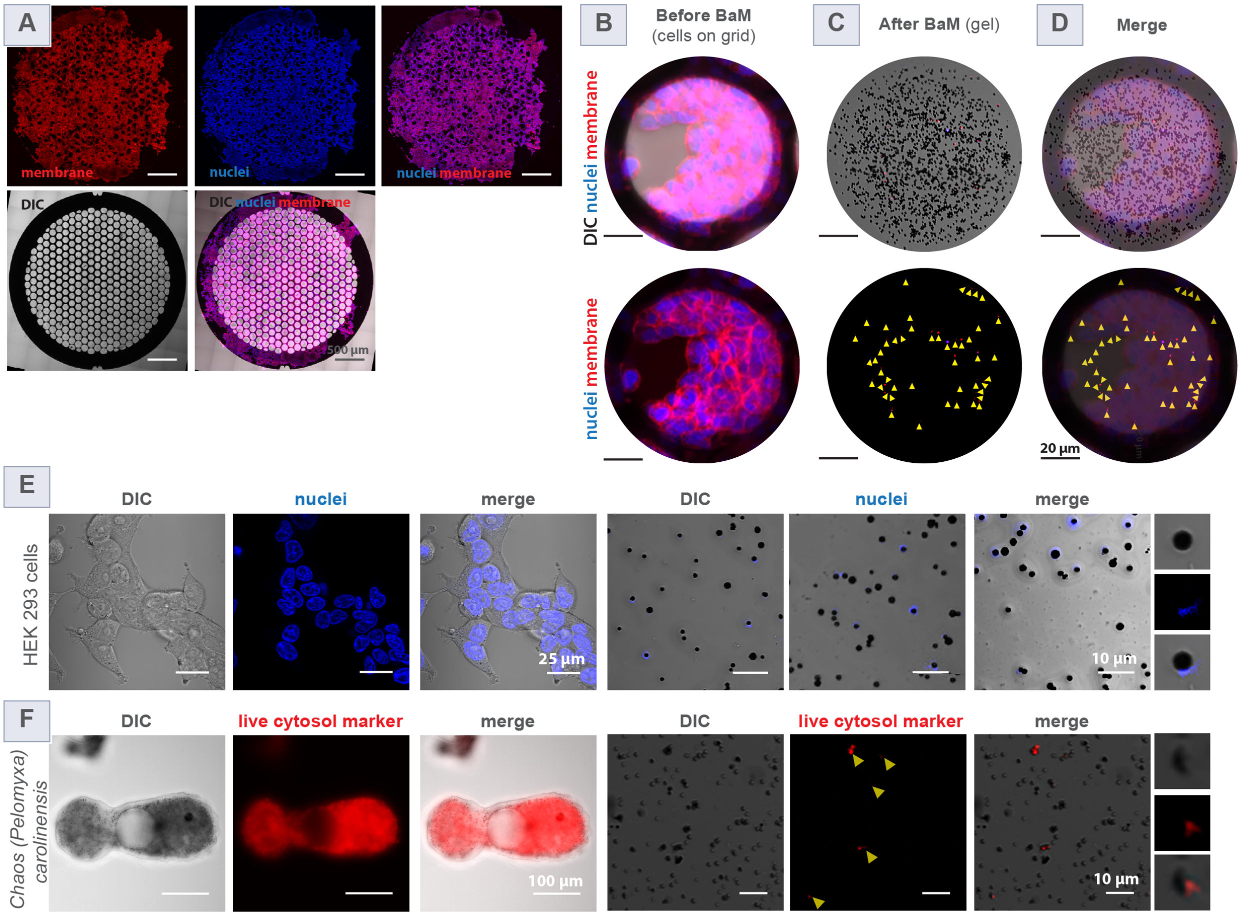


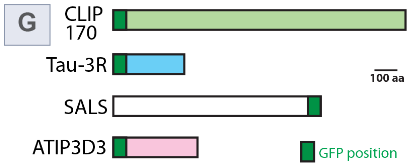


| Protein name | species | GFP position | size (aa) |
|---|---|---|---|
| Cytoplasmic Linker Protein of 170 kDa (CLIP 170) | mus musculus | N terminus | 1300 |
| Tau 3R | mus musculus | N terminus | 333 |
| Sarcomere length short (Sals) | drosophila melanogaster | C terminus | 933 |
| Angiotensin II type 2 (AT (2)) receptor-interacting protein 3 (ATIP3) Domain 3 | homo sapiens | N terminus | 396 |

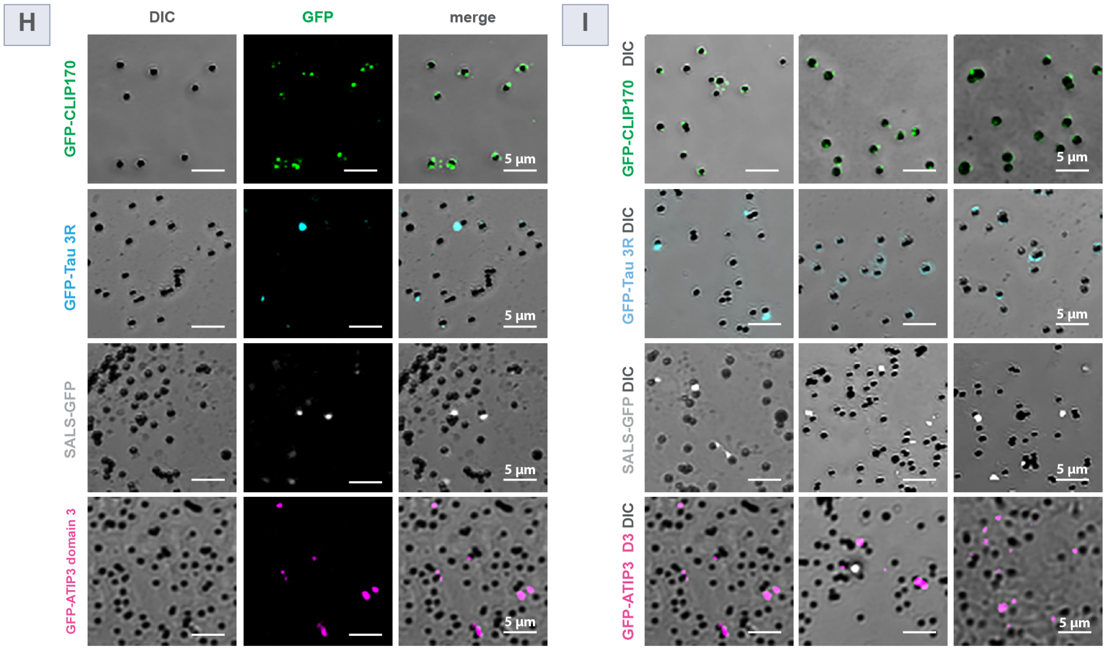




**Figure S3**: (Caption next page.)

**Figure S3**: **Supporting data for BaM of live cells and subsequent analysis using fluorescence microscopy (complementary to Fig. 3).** (A) DIC and fluorescence images of live HEK293 cells grown on an EM grid stained with Hoechst (nuclei, blue) and CellMask Orange (membrane, red), showing individual and merged channels. These images complement Figure 3B. (B-D) Magnified field of view of a single EM grid hole and corresponding BaM sampling images (related to Fig. 3E-G). (B) Pre-BaM fluorescence image of labeled live cells, shown with and without DIC. (C) Post-BaM fluorescence image of the gel showing embedded BaM particles, presented with and without DIC overlay. (D) Fluorescence overlay indicating successful collection of fluorescence signals by individual BaM particles. (E-F) Additional BaM experiments demonstrating the collection of nuclear material (Hoechst-stained) from HEK293 cells and cytoplasmic material from the non-model organism Chaos (Pelomyxa) carolinensis. (E) DIC, fluorescence, and merged images of HEK293 cells stained with Hoechst before BaM, and BaM particles post-BaM. (F) DIC, fluorescence, and merged images of Chaos (Pelomyxa) carolinensis stained with a live cytosolic marker before BaM, and particles after BaM. (G) Proteins used in this study that are known to form protein condensates or aggregates. Listed are protein names, species of origin, GFP tag position, and total size in amino acids. (H-I) Additional fields of view of BaM particles collecting GFP-tagged proteins from live cells: GFP-CLIP170 (green), GFP-Tau 3R (cyan), SALS-GFP (white), and GFP-ATIP3 domain 3 (magenta). These datasets complement Figure 3M.

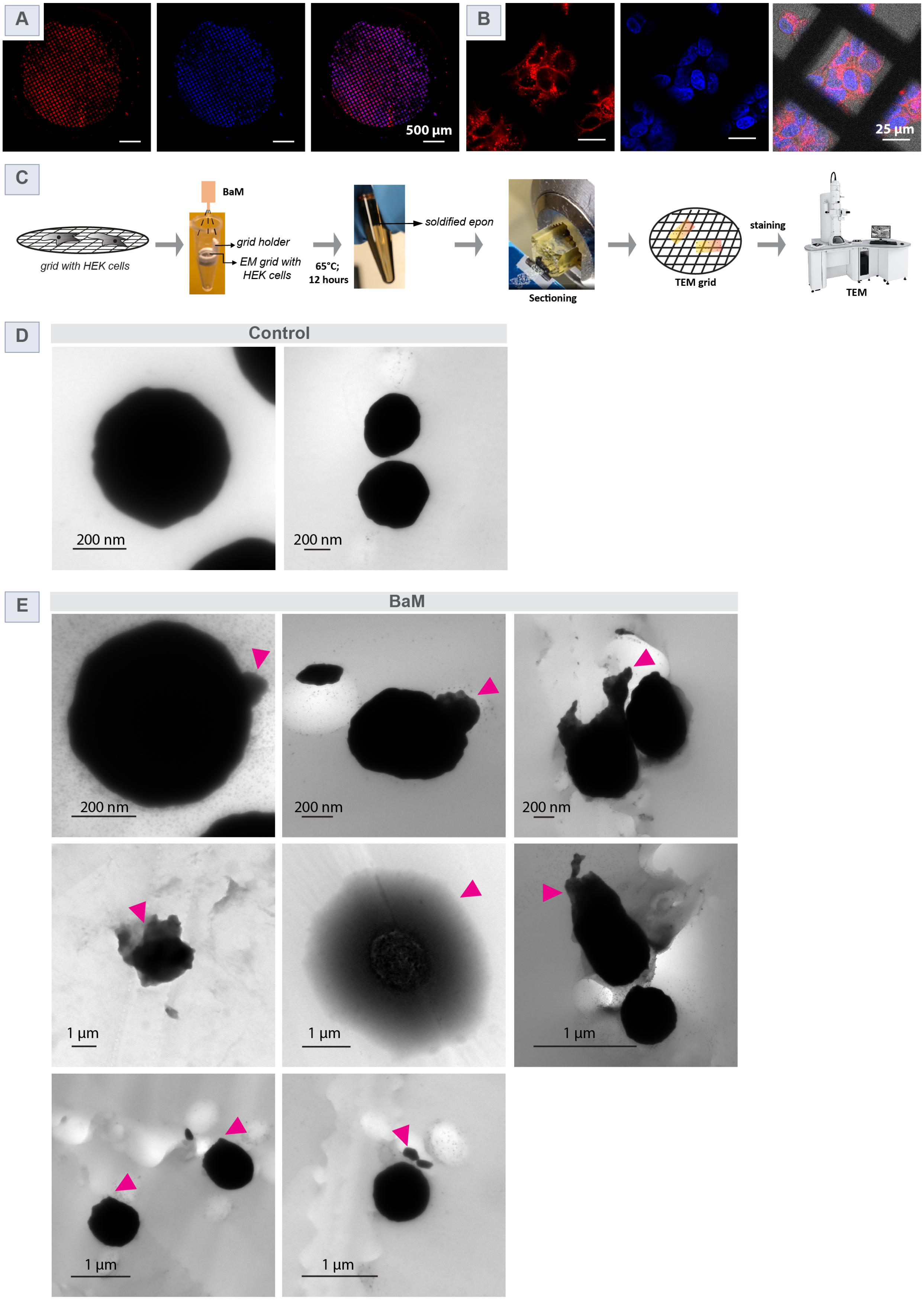




**Figure S4**: (Caption next page.)

**Figure S4**: **Transmission electron microscopy (TEM) of BaM particles carrying cellular materials.** Fluorescence images of live HEK293 cells cultured on a gold EM grid, stained with Hoechst (nuclei, blue) and CellMask Orange (membrane, red). (A) Full-grid view; (B) zoomed-in images showing individual cells on the grid surface (complement to Fig. 4D). (C) Schematic representation of the pipeline used to prepare BaM particles for TEM analysis. HEK293 cells were grown on a gold EM grid and placed in a custom grid holder inside a PCR tube filled with Epon resin. The grid was then bombarded with BaM particles (0.6-1 $\mu$m). Following bombardment, the EM grid and holder were carefully removed, and the resin was immediately transferred to 65 °C for polymerization and incubated for 12 hours. The solidified Epon resin block was sectioned using a Leica ultramicrotome, followed by staining and imaging with a transmission electron microscope. Additional details are provided in the Methods section. (D-E) Unprocessed transmission electron micrographs of BaM particles carrying cellular material. (D) BaM particles from the control experiment, in which an EM grid without cells was bombarded. (E) Electron micrographs of BaM particles bombarded through an EM grid with live HEK 293 cells. Pink arrowheads highlight various cellular materials associated with BaM particles, including cytoplasmic fragments and membranous structures.

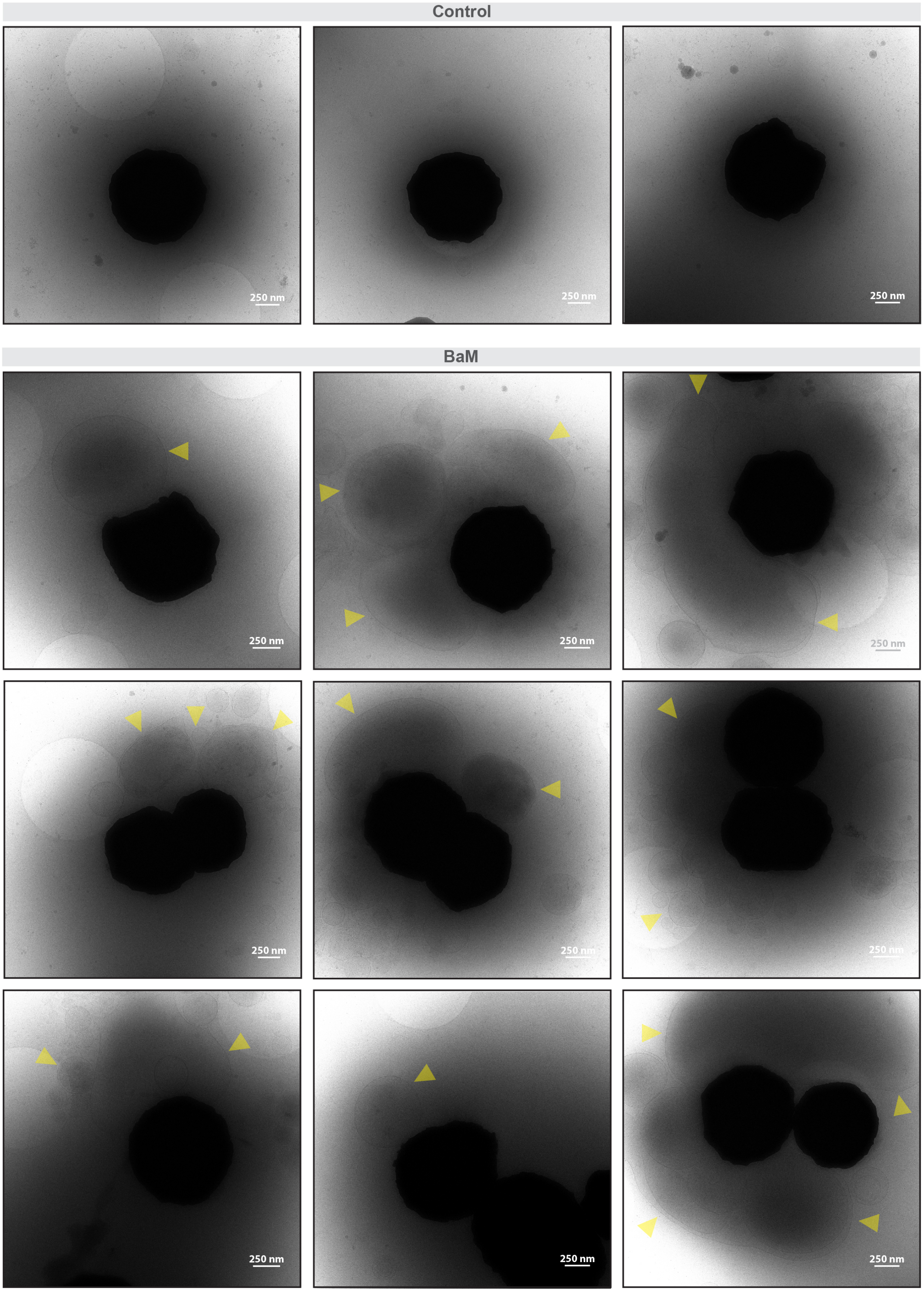




**Figure S5**: (Caption next page.)

**Figure S5**: **High-resolution cryo-electron micrographs reveal membrane-enclosed subcellular structures associated with BaM particles.** Control and BaM samples with 1 $\mu$m diameter gold microparticles imaged by Cryo-EM. The top panel (control) shows BaM particles without any associated cellular material. Bottom panels (BaM): BaM particles collected after bombardment through HEK293 cells exhibit electron-lucent structures surrounding or adjacent to the particles (yellow arrowheads), suggestive of membrane-like or vesicular material. These structures were absent in the control condition, supporting successful retrieval of subcellular content by BaM. Scale bar: 250 nm (all panels).

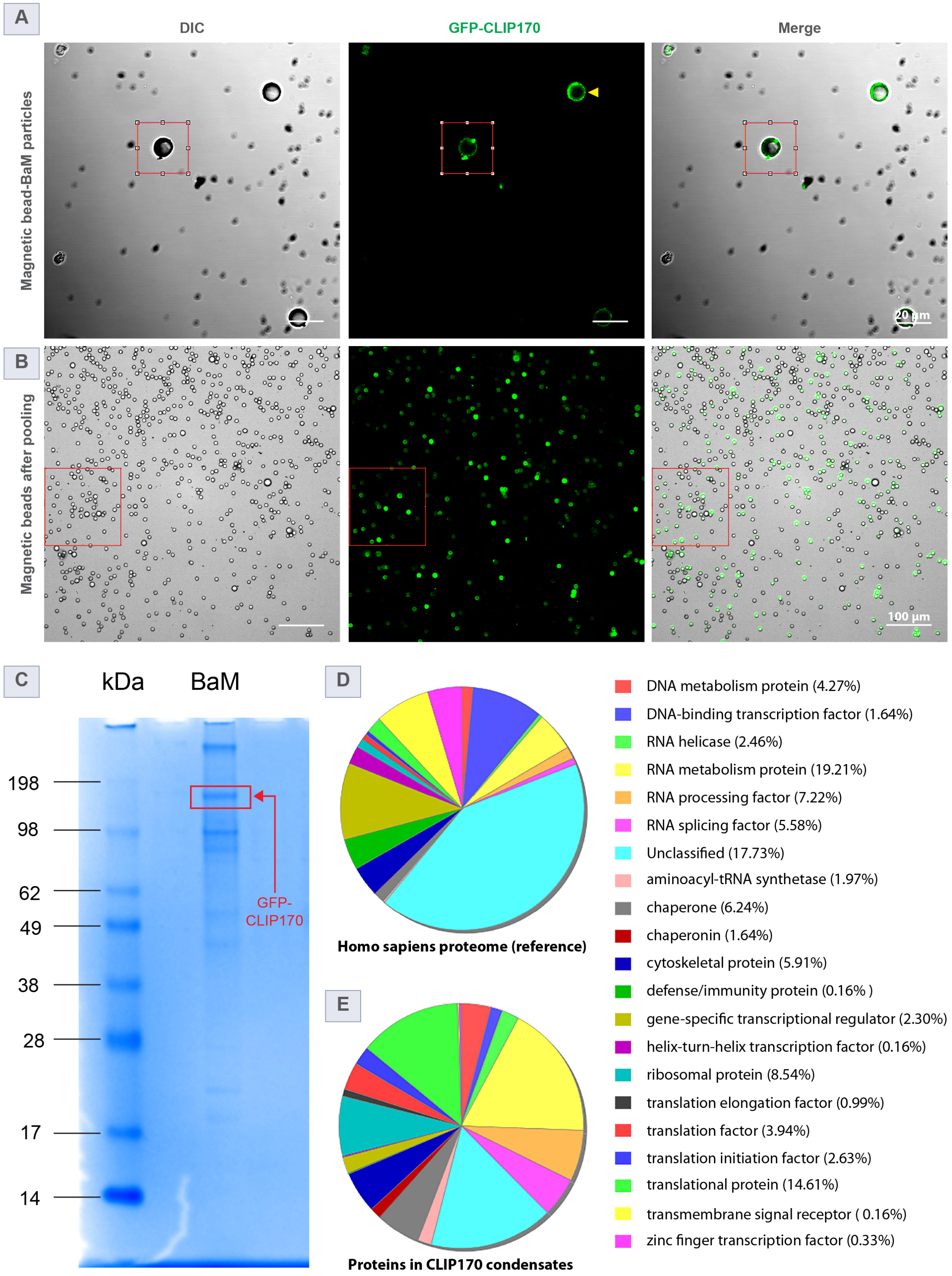




**Figure S6**: (Caption next page.)

**Figure S6**: **Supporting analysis of pooled GFP-CLIP170 condensates collected by BaM.** (A) Full field-of-view DIC, GFP fluorescence, and merged images showing magnetic beads (8 $\mu$m diameter) functionalized with anti-GFP antibodies. These beads are used to pool GFP-CLIP170 initially collected by BaM particles (see Fig. 5D). The red box highlights a region in Fig. 5D. Over time, we also observed the detachment of GFP-CLIP170 from gold particles and its subsequent association with anti-GFP–coated magnetic particles (indicated by a yellow arrowhead). This interaction and transfer from BaM particles to magnetic particles increases following the washing steps. (B) Fluorescence and DIC images showing a population of magnetic beads following pooling of GFP-CLIP170 condensates, prior to mass spectrometry analysis (see Fig. 5E). (C) Coomassie-stained SDS-PAGE gel showing total protein bound to magnetic beads from pooled BaM samples. The red box marks the band corresponding to GFP-tagged CLIP170. (D-E) Classification of proteins identified in the GFP-CLIP170 condensate using the PANTHER classification system (www.pantherdb.org). (D) Homo sapiens proteome as reference. (E) Protein composition of the CLIP170 condensate sample shows that 17.73% corresponds to unclassified proteins, with strong enrichment for RNA-binding and RNA metabolism-related categories, as well as translational proteins and chaperones. Percentages in the legend refer to the chart in (E) only.

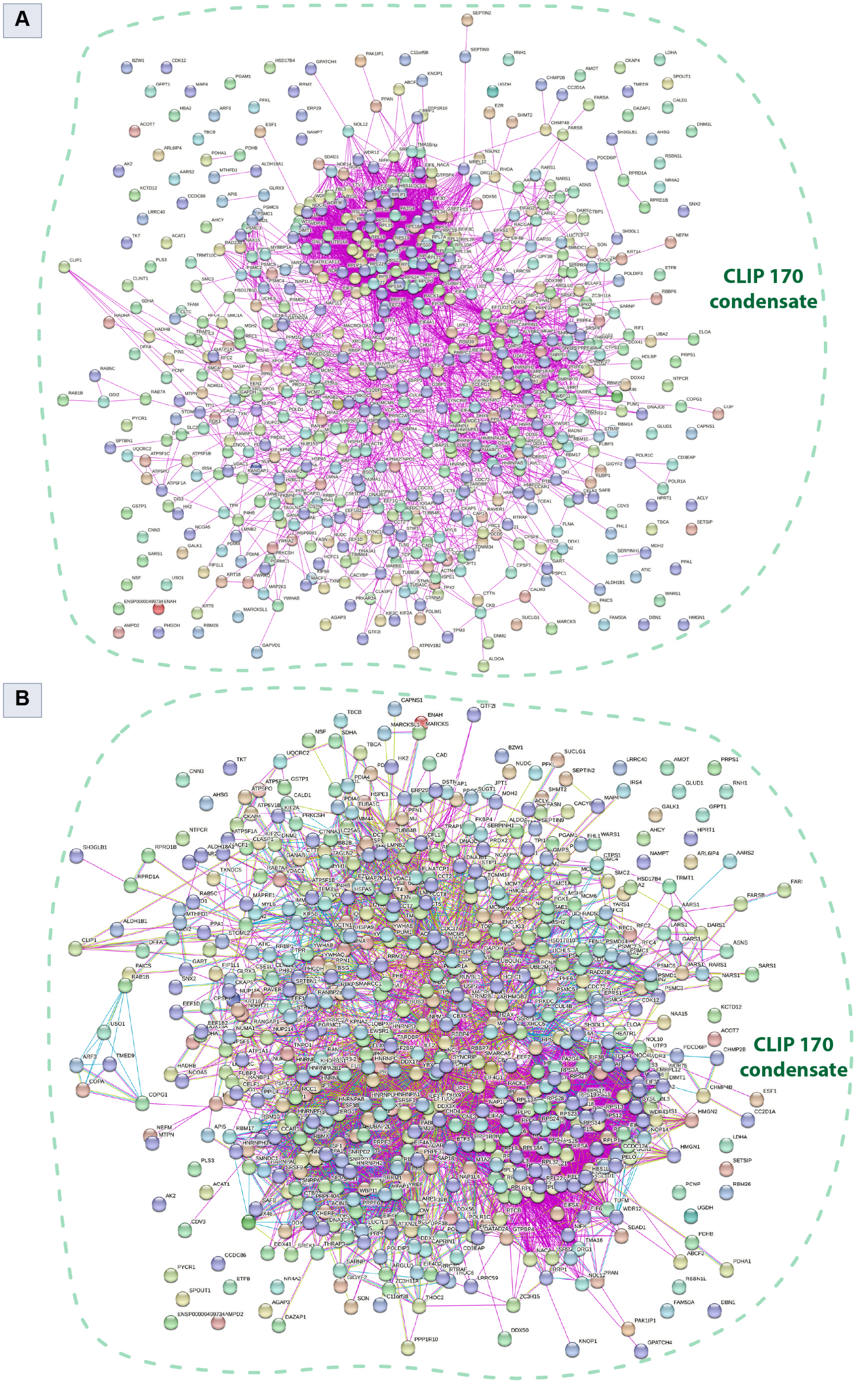




**Figure S7**: (Caption next page.)

**Figure S7**: **Protein-protein interaction (PPI) network analysis of proteins identified in CLIP170 condensate.** (A-B) Visualization of protein-protein interaction networks constructed using the STRING database for proteins identified in GFP-CLIP170 condensates (see Supplementary Table 2 for the full protein list). Each node represents a protein, and edges indicate predicted or known interactions. (A) Network showing only experimentally validated interactions (magenta lines). (B) Full STRING network showing both experimental and predicted associations. Edge types indicate: gene neighborhood (green), gene fusion (red), co-occurrence (blue), text mining (light green), co-expression (black), and protein homology (violet). The network reveals a complex and integrated landscape of RNA-binding, translation, and cytoskeletal proteins enriched in CLIP170 condensates.

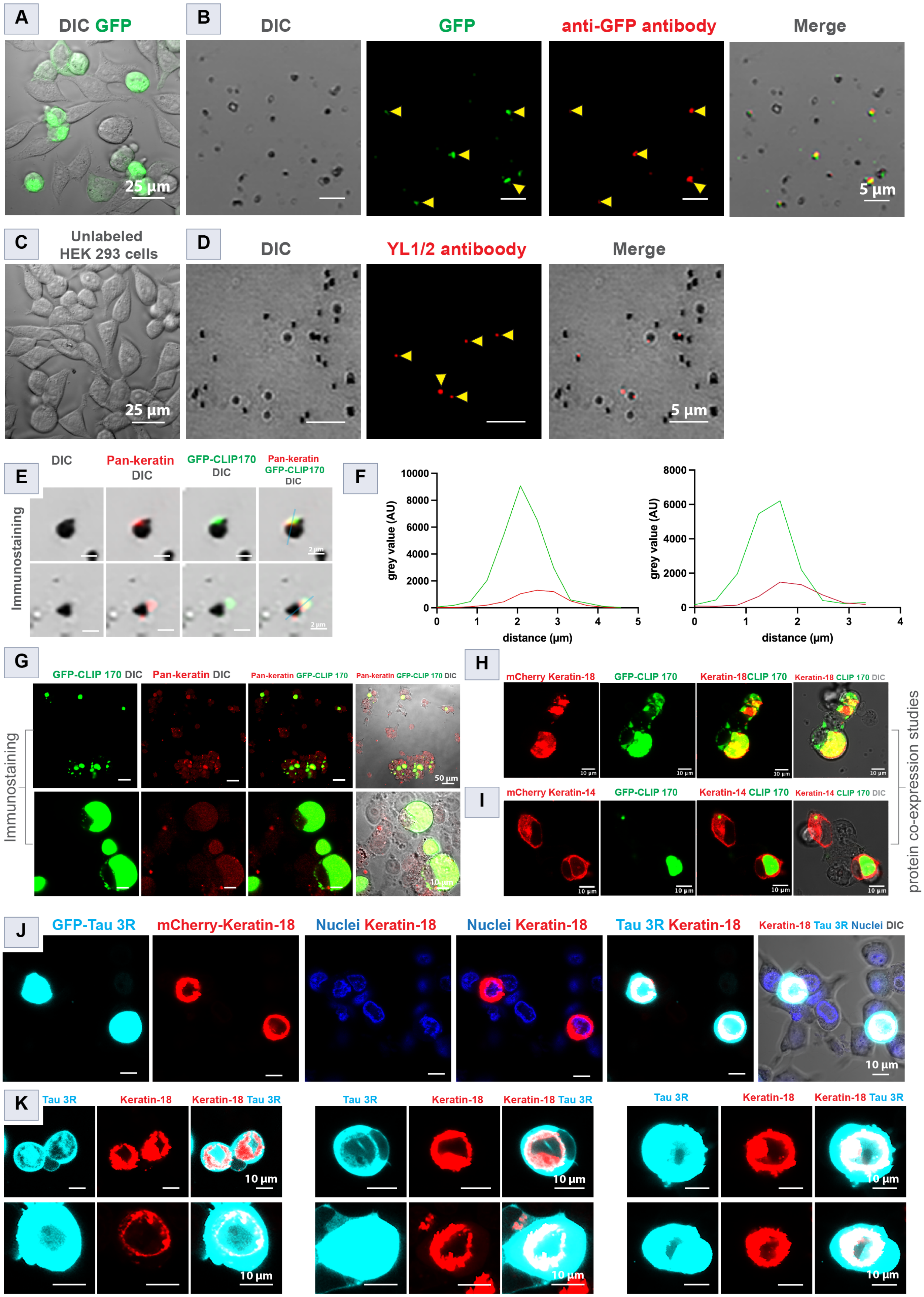




**Figure S8**: (Caption next page.)

**Figure S8**: **BaM particle-associated protein detection and co-localization analysis of CLIP170, Keratins, and Tau condensates (complementary to Fig. 3).** (A-C) Demonstration of immunostaining feasibility for cytosolic material on BaM particles. (A) Merged DIC and fluorescence image of HEK293 cells expressing GFP. (B) GFP-positive BaM particles (green) confirmed by anti-GFP antibody staining (red); yellow arrowheads indicate co-localization. (C) DIC image of unstained HEK293 cells (control). (D) Fluorescence and DIC images of BaM particles immunostained with YL1/2 antibody targeting tyrosinated tubulin; yellow arrowheads highlight immunopositive particles. (E) Immunostaining of BaM particles carrying GFP-tagged CLIP170 (green) with a pan-Keratin antibody (red), revealing co-localization on individual particles. (F) Line profile analysis of particles in (E) confirms spatial overlap of GFP-CLIP170 and pan-Keratin signals. (G) Pan-keratin staining of fixed HEK293 cells expressing GFP-CLIP170 shows strong co-localization with CLIP170 condensates. (H) Live-cell co-expression of GFP-CLIP170 (green) and mCherry-Keratin-18 (red) shows condensate-level co-localization; merged channels confirm spatial overlap. (I) Live-cell co-expression of GFP–CLIP170 (green) and mCherry–Keratin-14 (red) shows orthogonal localization, serving as a control. (J) Live-cell imaging of GFP-Tau 3R (cyan) and mCherry–Keratin-18 (red) in HEK293 cells, with DAPI-stained nuclei (blue). Merged images reveal a cage-like organization of Keratin-18 around Tau 3R condensates. (K) GFP-Tau 3R condensate (cyan)and mCherry-Keratin-18 (red) localization in multiple live single cells.

#### Supplementary Movie Captions

**See Movie S1:** https://drive.google.com/file/d/1BC8MsMcmZOaJW41qLphT2L5gEU5pormu/view?usp=drive_link

**Movie S1: High-speed imaging of BaM particles at the air-gel interface (related to Fig. 2G and Fig. S1H).** Representative time-lapse frames showing 5.1 $\mu$m BaM particles traversing the air-gel interface (1% agarose gel, 20 psi). Images were acquired using a high-speed camera (10× objective, 4788 frames per second). Still frames from this movie are shown in Fig. 2G and Fig. S1H. Time stamp, ms; scale bar, 200 $\mu$m.

**See Movie S2:** https://drive.google.com/file/d/1bs8PEUDmktbc6W_ZW2dVoBGoNEw8lx-v/view?usp=drive_link

**Movie S2: Z-stack video of BaM particles penetrating live HEK293 cells and carrying cellular material (related to Fig. 3I and Fig. S3B-D).** Z-stack imaging of 1 $\mu$m BaM particles penetrating live HEK293 cells stained with fluorescent markers (red: cell membrane; blue: nuclei). The video shows cellular material collected along the BaM particle trajectory. Panel A displays a merged view of DIC, membrane, and nuclear channels. Panel B shows the same field without the DIC channel, highlighting the fluorescent trail of cellular material behind the BaM particle.

**See Movie S3:** https://drive.google.com/file/d/1DMtDLSpVA2vfO5H8qpnkQose2oxOBq9O/view?usp=drive_link

**Movie S3: Cryo-EM tilt series of a BaM particle carrying membrane enclosed subcellular structures (corresponding to figure 4H-I).** Cryo-tomography of an individual BaM particle (500 nm radius) carrying cellular material. Panel A shows the raw file off the electron micrograph. Panel B shows a full 360° 3D reconstruction, with the BaM particle pseudo-colored in gold and the associated membrane-enclosed subcellular contents shown in gray.

**See Movie S4:** https://drive.google.com/file/d/1-2NCkSKEuQ-u5lOUin-g5TF1KqsxjCzG/view?usp=drive_link

**Movie S4: 3D visualization of CLIP170 condensates and Keratin localization in live HEK293 cells (corresponding to Fig. 5M–P and Fig. S8H).** A 3D reconstruction from confocal Z-stack images of live HEK293 cells expressing CLIP170 condensates and Keratin-18/14. CLIP170 is tagged with GFP (green), and Keratin-18/14 with mCherry (red). The video shows both individual and merged fluorescence channels, visualizing the spatial organization and co-localization of CLIP170 condensates with Keratin proteins.

**See Movie S5:** `https://drive.google.com/file/d/18UVU0iJ2Af6485cjEDHujTSmsZuc-rRi/view?usp=sharing`

**Movie S5: 3D visualization of Tau-3R condensates and Keratin-18 localization in live HEK293 cells (corresponding to Fig. 5L and Fig. S8J).** A 3D reconstruction from confocal Z-stack images of live HEK293 cells expressing Tau-3R condensates and Keratin-18. Tau-3R is tagged with GFP (cyan), and Keratin-18 with mCherry (red). The upper panel shows the full Z-stack of the HEK293 cell, while the bottom panel presents a cross-section, highlighting the spatial co-localization of Tau-3R condensates with Keratin-18. Panel A (last column) shows individual Z-planes spanning from one end of the cell to the other.

**Table S1: Physical regime of Ballistic Microscopy.**

**Table S2: List of proteins identified in pooled GFP-CLIP170 condensates using BaM and mass spectrometry.** A total of 641 proteins were detected in the GFP-CLIP170 condensates pooled with GFP-coated magnetic beads from BaM particles (complementary to Fig. 5F and Fig. S7A,B).

**Table S2. List of proteins identified in pooled GFP-CLIP170 condensates using Ballistic Microscopy and mass spectrometry.** A total of 641 proteins were detected in the GFP-CLIP170 condensates pooled with GFP-coated magnetic beads from BaM particles (complementary to Fig. 5G and Fig. S7A,B).

| Sl no. | Protein name | Protein ID |
|---|---|---|
| 1 | CAP-Gly domain-containing linker protein 1 isoform b | P30622 |
| 2 | Pyruvate kinase | A0A8V8TNX9 |
| 3 | Elongation factor 2 | P13639 |
| 4 | D-3-phosphoglycerate dehydrogenase | O43175 |
| 5 | Actin, cytoplasmic 1 | P60709 |
| 6 | Filamin-A | P21333 |
| 7 | Fatty acid synthase | A0A0U1RQF0 |
| 8 | T-complex protein 1 subunit beta | P78371 |
| 9 | Talin-1 | Q9Y490 |
| 10 | ATP-dependent RNA helicase A | Q08211 |
| 11 | Probable ATP-dependent RNA helicase DDX5 | P17844 |
| 12 | C-1-tetrahydrofolate synthase, cytoplasmic | P11586 |
| 13 | Isoform 2 of Nuclear mitotic apparatus protein 1 | Q14980 |
| 14 | Transcription intermediary factor 1-beta | Q13263 |
| 15 | Isoform 2 of Heterogeneous nuclear ribonucleoprotein K | P61978 |
| 16 | Heterogeneous nuclear ribonucleoprotein U | Q00839 |
| 17 | Cytoplasmic dynein 1 heavy chain 1 | Q14204 |
| 18 | Bifunctional glutamate/proline--tRNA ligase | P07814 |
| 19 | Isoform 2 of RNA-binding protein 39 | Q14498 |
| 20 | ATP synthase subunit alpha, mitochondrial | P25705 |
| 21 | Nucleophosmin | P06748 |
| 22 | ATP-dependent RNA helicase DDX3X | O00571 |
| 23 | Isoform 4 of DNA topoisomerase 2-alpha | P11388 |
| 24 | Nucleolin | P19338 |
| 25 | Ubiquitin-associated protein 2-like | Q14157 |
| 26 | Ubiquitin-like modifier-activating enzyme 1 | P22314 |
| 27 | Stress-70 protein, mitochondrial | P38646 |
| 28 | ATP synthase subunit beta, mitochondrial | P06576 |
| 29 | Matrin-3 | P43243 |
| 30 | T-complex protein 1 subunit epsilon | P48643 |
| 31 | Isoform 8 of Eukaryotic translation initiation factor 4 gamma 1 | Q04637 |
| 32 | Elongation factor Tu, mitochondrial | P49411 |
| 33 | DNA-dependent protein kinase catalytic subunit | P78527 |
| 34 | Endoplasmic reticulum chaperone BiP | P11021 |
| 35 | T-complex protein 1 subunit gamma | P49368 |
| 36 | T-complex protein 1 subunit eta | Q99832 |
| 37 | Heat shock protein HSP 90-alpha | P07900 |
| 38 | T-complex protein 1 subunit theta | P50990 |
| 39 | T-complex protein 1 subunit delta | P50991 |
| 40 | Far upstream element-binding protein 1 | Q96AE4 |

| | | |
|---|---|---|
| 41 | Stalled ribosome sensor GCN1 | Q92616 |
| 42 | Peroxiredoxin-1 | Q06830 |
| 43 | Transgelin-2 | P37802 |
| 44 | RNA helicase | A0A0C4DG89 |
| 45 | Heterogeneous nuclear ribonucleoproteins A2/B1 | P22626 |
| 46 | Probable ATP-dependent RNA helicase DDX17 | Q92841 |
| 47 | T-complex protein 1 subunit alpha | P17987 |
| 48 | Nuclear autoantigenic sperm protein | P49321 |
| 49 | Acetyl-CoA acetyltransferase, mitochondrial | P24752 |
| 50 | Inosine-5'-monophosphate dehydrogenase | A0A994J749 |
| 51 | Fructose-bisphosphate aldolase A | P04075 |
| 52 | Alanine--tRNA ligase, cytoplasmic | P49588 |
| 53 | X-ray repair cross-complementing protein 6 | P12956 |
| 54 | T-complex protein 1 subunit zeta | P40227 |
| 55 | RuvB-like 2 | Q9Y230 |
| 56 | Pre-mRNA-processing-splicing factor 8 | Q6P2Q9 |
| 57 | Stress-induced-phosphoprotein 1 | P31948 |
| 58 | Clathrin heavy chain | A0A087WVQ6 |
| 59 | Alpha-enolase | P06733 |
| 60 | Angiomotin | Q4VCS5 |
| 61 | Eukaryotic initiation factor 4A-I | P60842 |
| 62 | Splicing factor 3B subunit 2 | Q13435 |
| 63 | Small ribosomal subunit protein eS4, X isoform | P62701 |
| 64 | Heterogeneous nuclear ribonucleoprotein H | P31943 |
| 65 | Heterogeneous nuclear ribonucleoprotein A1 | P09651 |
| 66 | X-ray repair cross-complementing protein 5 | P13010 |
| 67 | Glycine--tRNA ligase | P41250 |
| 68 | Signal recognition particle subunit SRP68 | Q9UHB9 |
| 69 | Exosome complex exonuclease RRP44 | Q9Y2L1 |
| 70 | ATP-dependent RNA helicase DDX42 | Q86XP3 |
| 71 | Polyadenylate-binding protein 1 | P11940 |
| 72 | Transcription factor BTF3 | P20290 |
| 73 | Regulator of chromosome condensation | P18754 |
| 74 | Myosin-10 | P35580 |
| 75 | Histone H2B type 1-O | P23527 |
| 76 | Mediator of DNA damage checkpoint protein 1 | Q14676 |
| 77 | Lamina-associated polypeptide 2, isoforms beta/gamma | P42167 |
| 78 | Guanine nucleotide-binding protein-like 3 | Q9BVP2 |
| 79 | Bifunctional phosphoribosylaminoimidazole carboxylase/phosphoribosylaminoimidazole succinocarboxamide synthetase | P22234 |
| 80 | Elongation factor 1-gamma | P26641 |
| 81 | Heterogeneous nuclear ribonucleoprotein L | P14866 |
| 82 | Polypyrimidine tract-binding protein 1 | P26599 |
| 83 | CTP synthase 1 | P17812 |

| | | |
|---|---|---|
| 84 | DNA replication licensing factor MCM7 | P33993 |
| 85 | Glyceraldehyde-3-phosphate dehydrogenase | P04406 |
| 86 | Transitional endoplasmic reticulum ATPase | P55072 |
| 87 | Creatine kinase B-type | P12277 |
| 88 | Nuclear receptor subfamily 4 group A member 2 | P43354 |
| 89 | Heat shock protein 75 kDa, mitochondrial | Q12931 |
| 90 | Plastin-3 | P13797 |
| 91 | Lamin-B1 | P20700 |
| 92 | Phosphoglycerate kinase 1 | P00558 |
| 93 | Microtubule-associated protein | E7EVA0 |
| 94 | Small ribosomal subunit protein eS17 | P08708 |
| 95 | Isoform 2 of Splicing factor U2AF 65 kDa subunit | P26368 |
| 96 | Isoform 3 of Poly(rC)-binding protein 2 | Q15366 |
| 97 | Small ribosomal subunit protein eS7 | P62081 |
| 98 | 116 kDa U5 small nuclear ribonucleoprotein component | Q15029 |
| 99 | Triosephosphate isomerase | P60174 |
| 100 | Large ribosomal subunit protein uL3 | P39023 |
| 101 | RuvB-like 1 | Q9Y265 |
| 102 | ATP-dependent RNA helicase DDX1 | Q92499 |
| 103 | Peptidyl-prolyl cis-trans isomerase A | P62937 |
| 104 | Large ribosomal subunit protein P2 | P05387 |
| 105 | Bcl-2-associated transcription factor 1 | Q9NYF8 |
| 106 | Insulin-like growth factor 2 mRNA-binding protein 1 | Q9NZI8 |
| 107 | Regulator of nonsense transcripts 1 | Q92900 |
| 108 | Complement component 1 Q subcomponent-binding protein, mitochondrial | Q07021 |
| 109 | Sodium/potassium-transporting ATPase subunit alpha-1 | P05023 |
| 110 | Nucleolar protein 58 | Q9Y2X3 |
| 111 | Pre-mRNA-processing factor 6 | O94906 |
| 112 | Protein disulfide-isomerase | P07237 |
| 113 | Cytoskeleton-associated protein 4 | Q07065 |
| 114 | RNA cytosine C(5)-methyltransferase NSUN2 | Q08J23 |
| 115 | Eukaryotic translation initiation factor 5B | A0A087WUT6 |
| 116 | Staphylococcal nuclease domain-containing protein 1 | Q7KZF4 |
| 117 | Structural maintenance of chromosomes protein 1A | Q14683 |
| 118 | Trifunctional purine biosynthetic protein adenosine-3 | P22102 |
| 119 | RNA-binding protein 14 | Q96PK6 |
| 120 | Far upstream element-binding protein 3 | Q96I24 |
| 121 | Heterogeneous nuclear ribonucleoprotein A3 | A0A7I2V4G0 |
| 122 | Cytoskeleton-associated protein 5 | Q14008 |
| 123 | 14-3-3 protein zeta/delta | P63104 |
| 124 | Endoplasmin | P14625 |
| 125 | Vigilin | Q00341 |
| 126 | Isoform 4 of Protein PRRC2C | Q9Y520 |
| 127 | Scaffold attachment factor B1 | Q15424 |

| | | |
|---|---|---|
| 128 | Splicing factor 3A subunit 1 | Q15459 |
| 129 | Interleukin enhancer-binding factor 3 | Q12906 |
| 130 | Leucine--tRNA ligase, cytoplasmic | Q9P2J5 |
| 131 | Isoform 2 of Protein enabled homolog | Q8N8S7 |
| 132 | Apoptosis inhibitor 5 | Q9BZZ5 |
| 133 | Isoform 2 of Elongation factor 1-delta | P29692 |
| 134 | Flap endonuclease 1 | P39748 |
| 135 | Ribosomal RNA processing protein 1 homolog B | Q14684 |
| 136 | Leucine-rich PPR motif-containing protein, mitochondrial | P42704 |
| 137 | Importin subunit alpha-1 | P52292 |
| 138 | Multifunctional protein CAD | P27708 |
| 139 | RNA-binding motif protein, X chromosome | P38159 |
| 140 | Ras GTPase-activating-like protein IQGAP1 | P46940 |
| 141 | Cell cycle and apoptosis regulator protein 2 | Q8N163 |
| 142 | Heterogeneous nuclear ribonucleoprotein H3 | P31942 |
| 143 | Splicing factor 1 | H7C561 |
| 144 | Polymerase delta-interacting protein 3 | Q9BY77 |
| 145 | General vesicular transport factor p115 | O60763 |
| 146 | Malate dehydrogenase, mitochondrial | P40926 |
| 147 | Spliceosome RNA helicase DDX39B | Q13838 |
| 148 | Peptidyl-prolyl cis-trans isomerase FKBP4 | Q02790 |
| 149 | Small ribosomal subunit protein eS6 | P62753 |
| 150 | Cofilin-1 | P23528 |
| 151 | serine--tRNA ligase | Q5T5C7 |
| 152 | SNW domain-containing protein 1 | Q13573 |
| 153 | Large ribosomal subunit protein uL4 | P36578 |
| 154 | Double-stranded RNA-specific adenosine deaminase | P55265 |
| 155 | Large ribosomal subunit protein uL11 | P30050 |
| 156 | Eukaryotic translation initiation factor 3 subunit D | O15371 |
| 157 | Isoform 2 of Tropomyosin alpha-3 chain | P06753 |
| 158 | SERPINE1 mRNA-binding protein 1 | Q8NC51 |
| 159 | Heterogeneous nuclear ribonucleoprotein C | B4DY08 |
| 160 | Kinesin-1 heavy chain | P33176 |
| 161 | Delta-1-pyrroline-5-carboxylate synthase | P54886 |
| 162 | Nucleoprotein TPR | P12270 |
| 163 | DNA replication licensing factor MCM3 | P25205 |
| 164 | Y-box-binding protein 1 | P67809 |
| 165 | 26S proteasome regulatory subunit 10B | P62333 |
| 166 | Programmed cell death 6-interacting protein | Q8WUM4 |
| 167 | Hypoxanthine-guanine phosphoribosyltransferase | P00492 |
| 168 | ADP-ribosylation factor 3 | P61204 |
| 169 | Eukaryotic translation initiation factor 4 gamma 2 | P78344 |
| 170 | Serine/threonine-protein phosphatase 2A 65 kDa regulatory subunit A alpha isoform | P30153 |
| 171 | RNA-splicing ligase RtcB homolog | Q9Y3I0 |

| | | |
|---|---|---|
| 172 | HEAT repeat-containing protein 1 | Q9H583 |
| 173 | ATP-citrate synthase | P53396 |
| 174 | Profilin-1 | P07737 |
| 175 | WD repeat-containing protein 36 | Q8NI36 |
| 176 | Serine hydroxymethyltransferase, mitochondrial | P34897 |
| 177 | Asparagine synthetase [glutamine-hydrolyzing] | P08243 |
| 178 | Tubulin beta-4B chain | P68371 |
| 179 | GTP-binding protein 4 | Q9BZE4 |
| 180 | Aspartate--tRNA ligase, cytoplasmic | P14868 |
| 181 | Probable ATP-dependent RNA helicase DDX41 | Q9UJV9 |
| 182 | 14-3-3 protein epsilon | P62258 |
| 183 | Importin subunit beta-1 | Q14974 |
| 184 | Eukaryotic translation initiation factor 5A-1 | P63241 |
| 185 | TAR DNA-binding protein 43 | Q13148 |
| 186 | Small ribosomal subunit protein eS8 | P62241 |
| 187 | Serrate RNA effector molecule homolog | Q9BXP5 |
| 188 | Cleavage and polyadenylation specificity factor subunit 6 | Q16630 |
| 189 | Melanoma-associated antigen D2 | Q9UNF1 |
| 190 | DnaJ homolog subfamily C member 7 | Q99615 |
| 191 | Ras GTPase-activating protein-binding protein 1 | Q13283 |
| 192 | L-lactate dehydrogenase B chain | P07195 |
| 193 | Ribonuclease inhibitor | P13489 |
| 194 | PHD finger protein 6 | Q8IWS0 |
| 195 | Small ribosomal subunit protein uS8 | P62244 |
| 196 | Heat shock protein 105 kDa | Q92598 |
| 197 | Probable 28S rRNA (cytosine(4447)-C(5))-methyltransferase | P46087 |
| 198 | Isoform 2 of Splicing regulatory glutamine/lysine-rich protein 1 | Q8WXA9 |
| 199 | Transducin beta-like protein 3 | Q12788 |
| 200 | Ubiquitin-ribosomal protein eS31 fusion protein | P62979 |
| 201 | Tyrosine--tRNA ligase, cytoplasmic | P54577 |
| 202 | Structural maintenance of chromosomes protein 4 | Q9NTJ3 |
| 203 | Cleavage and polyadenylation specificity factor subunit 7 | Q8N684 |
| 204 | Prohibitin 1 | P35232 |
| 205 | Trifunctional enzyme subunit alpha, mitochondrial | P40939 |
| 206 | Isoform 3 of Dynamin-2 | P50570 |
| 207 | FACT complex subunit SSRP1 | Q08945 |
| 208 | Large ribosomal subunit protein uL10 | P05388 |
| 209 | Eukaryotic translation initiation factor 2 subunit 3 | P41091 |
| 210 | Pyruvate dehydrogenase E1 component subunit beta, mitochondrial | P11177 |
| 211 | Cyclin-dependent kinase 12 | Q9NYV4 |
| 212 | Calponin-3 | Q15417 |
| 213 | SAP domain-containing ribonucleoprotein | P82979 |
| 214 | Calnexin | A0A7P0TAE9 |
| 215 | Proliferation-associated protein 2G4 | Q9UQ80 |
| 216 | Glutamine--fructose-6-phosphate aminotransferase [isomerizing] 1 | Q06210 |

| | | |
|---|---|---|
| 217 | Protein NDRG1 | Q92597 |
| 218 | Myosin light chain 6 | G3V1V0 |
| 219 | 26S proteasome regulatory subunit 6A | P17980 |
| 220 | Signal recognition particle subunit SRP72 | O76094 |
| 221 | RRP12-like protein | Q5JTH9 |
| 222 | 5'-3' exoribonuclease 2 | Q9H0D6 |
| 223 | Eukaryotic initiation factor 4A-III | P38919 |
| 224 | Apoptotic chromatin condensation inducer in the nucleus | Q9UKV3 |
| 225 | THO complex subunit 2 | Q8NI27 |
| 226 | Host cell factor 1 | P51610 |
| 227 | Exportin-1 | O14980 |
| 228 | DNA replication licensing factor MCM5 | P33992 |
| 229 | 26S proteasome regulatory subunit 7 | P35998 |
| 230 | Replication factor C subunit 1 | P35251 |
| 231 | Inorganic pyrophosphatase | Q15181 |
| 232 | DNA-directed RNA polymerase I subunit RPA34 | O15446 |
| 233 | General transcription factor II-I | P78347 |
| 234 | Heterogeneous nuclear ribonucleoprotein D0 | Q14103 |
| 235 | DNA replication licensing factor MCM4 | P33991 |
| 236 | Arginine--tRNA ligase, cytoplasmic | P54136 |
| 237 | Galactokinase | P51570 |
| 238 | Isoform 2 of RNA-binding protein 26 | Q5T8P6 |
| 239 | KH domain-containing, RNA-binding, signal transduction-associated protein 1 | Q07666 |
| 240 | Isoform 2 of Heterogeneous nuclear ribonucleoprotein A/B | Q99729 |
| 241 | Exportin-2 | P55060 |
| 242 | Probable ATP-dependent RNA helicase DDX6 | P26196 |
| 243 | Chromobox protein homolog 3 | Q13185 |
| 244 | Isoleucine--tRNA ligase, cytoplasmic | P41252 |
| 245 | UV excision repair protein RAD23 homolog B | P54727 |
| 246 | Replication factor C subunit 4 | P35249 |
| 247 | ELAV-like protein 1 | Q15717 |
| 248 | Chromodomain-helicase-DNA-binding protein 4 | Q14839 |
| 249 | Prohibitin-2 | Q99623 |
| 250 | Heterogeneous nuclear ribonucleoprotein F | P52597 |
| 251 | Small ribosomal subunit protein eS10 | P46783 |
| 252 | Kinesin-like protein KIF2A | O00139 |
| 253 | Serpin H1 | P50454 |
| 254 | Src substrate cortactin | Q14247 |
| 255 | Something about silencing protein 10 | Q9NQZ2 |
| 256 | SWI/SNF-related matrix-associated actin-dependent regulator of chromatin subfamily A member 5 | O60264 |
| 257 | Insulin receptor substrate 4 | O14654 |
| 258 | G-patch domain and KOW motifs-containing protein | Q92917 |
| 259 | Large ribosomal subunit protein eL8 | P62424 |

| | | |
|---|---|---|
| 260 | Hsc70-interacting protein | P50502 |
| 261 | Replication protein A 70 kDa DNA-binding subunit | P27694 |
| 262 | Large ribosomal subunit protein eL6 | Q02878 |
| 263 | 14-3-3 protein beta/alpha | P31946 |
| 264 | Vinculin | P18206 |
| 265 | Enhanced Green Fluorescent Protein | 1373325 |
| 266 | BTB/POZ domain-containing protein KCTD12 | Q96CX2 |
| 267 | Developmentally-regulated GTP-binding protein 1 | Q9Y295 |
| 268 | MARCKS-related protein | P49006 |
| 269 | DnaJ homolog subfamily B member 1 | P25685 |
| 270 | Cytoplasmic linker associated protein 1 | F8WA11 |
| 271 | Neurofilament medium polypeptide | P07197 |
| 272 | Cold shock domain containing E1 | A0A8I5KV85 |
| 273 | Thyroid hormone receptor-associated protein 3 | Q9Y2W1 |
| 274 | THO complex subunit 4 | Q86V81 |
| 275 | Small ribosomal subunit protein uS5 | P15880 |
| 276 | Keratin, type I cytoskeletal 14 | P02533 |
| 277 | ATP-dependent RNA helicase DDX19A | Q9NUU7 |
| 278 | Adenylate kinase 2, mitochondrial | P54819 |
| 279 | Nascent polypeptide-associated complex subunit alpha, muscle-specific form | E9PAV3 |
| 280 | Calcyclin-binding protein | Q9HB71 |
| 281 | Ribonucleoprotein, PTB binding 1 | A0A087WZ13 |
| 282 | MKI67 FHA domain-interacting nucleolar phosphoprotein | Q9BYG3 |
| 283 | Protein disulfide-isomerase A6 | Q15084 |
| 284 | U3 small nucleolar RNA-associated protein 14 homolog A | Q9BVJ6 |
| 285 | Erythrocyte membrane protein band 4.1 like 2 | A0A994J5B1 |
| 286 | Nucleolar complex protein 4 homolog | Q9BVI4 |
| 287 | La-related protein 1 | Q6PKG0 |
| 288 | Heterogeneous nuclear ribonucleoprotein A0 | Q13151 |
| 289 | Paraspeckle component 1 | Q8WXF1 |
| 290 | Leucine-rich repeat-containing protein 40 | Q9H9A6 |
| 291 | DNA-directed RNA polymerase I subunit RPA1 | O95602 |
| 292 | Large ribosomal subunit protein uL6 | P32969 |
| 293 | WW domain-binding protein 11 | Q9Y2W2 |
| 294 | 26S proteasome non-ATPase regulatory subunit 4 | P55036 |
| 295 | Protein CDV3 homolog | Q9UKY7 |
| 296 | Isoform 4 of Heterogeneous nuclear ribonucleoprotein Q | O60506 |
| 297 | Coiled-coil domain-containing protein 86 | Q9H6F5 |
| 298 | Ribose-phosphate pyrophosphokinase 1 | P60891 |
| 299 | Ran-specific GTPase-activating protein | P43487 |
| 300 | Thioredoxin | P10599 |
| 301 | 26S proteasome regulatory subunit 6B | P43686 |
| 302 | Eukaryotic translation initiation factor 3 subunit C | Q99613 |
| 303 | Isoform 4 of Small ribosomal subunit protein eS24 | P62847 |

| | | |
|---|---|---|
| 304 | DNA repair protein RAD50 | Q92878 |
| 305 | Luc7-like protein 3 | O95232 |
| 306 | Ras-related protein Rab-1B | Q9H0U4 |
| 307 | RNA transcription, translation and transport factor protein | Q9Y224 |
| 308 | Small ribosomal subunit protein RACK1 | P63244 |
| 309 | RNA-binding protein 15 | Q96T37 |
| 310 | Hexokinase-2 | P52789 |
| 311 | U1 small nuclear ribonucleoprotein 70 kDa | P08621 |
| 312 | Translation machinery-associated protein 16 | Q96EY4 |
| 313 | H1.10 linker histone | A0A994J4R3 |
| 314 | Arf-GAP with GTPase, ANK repeat and PH domain-containing protein 3 | Q96P47 |
| 315 | Interleukin enhancer-binding factor 2 | Q12905 |
| 316 | Protein disulfide-isomerase A3 | P30101 |
| 317 | L-lactate dehydrogenase A chain | P00338 |
| 318 | Cytoplasmic linker associated protein 2 | A0A804HJG7 |
| 319 | Large ribosomal subunit protein eL22 | P35268 |
| 320 | Coatomer subunit alpha | P53621 |
| 321 | Transketolase | P29401 |
| 322 | Microtubule-associated protein RP/EB family member 1 | Q15691 |
| 323 | Mitotic checkpoint protein BUB3 | O43684 |
| 324 | Mitochondrial import receptor subunit TOM34 | Q15785 |
| 325 | Septin-2 | Q15019 |
| 326 | Pyrroline-5-carboxylate reductase 1, mitochondrial | P32322 |
| 327 | Large ribosomal subunit protein uL16 | P27635 |
| 328 | Protein SON | P18583 |
| 329 | High mobility group protein B1 | P09429 |
| 330 | Probable ATP-dependent RNA helicase DDX56 | Q9NY93 |
| 331 | Cancer-related nucleoside-triphosphatase | Q9BSD7 |
| 332 | Nucleosome assembly protein 1-like 1 | P55209 |
| 333 | Calcium homeostasis endoplasmic reticulum protein | Q8IWX8 |
| 334 | Core histone macro-H2A.1 | O75367 |
| 335 | SUMO-activating enzyme subunit 2 | Q9UBT2 |
| 336 | Alpha-actinin-4 | O43707 |
| 337 | Suppressor of SWI4 1 homolog | Q9NQ55 |
| 338 | Keratin, type II cytoskeletal 8 | P05787 |
| 339 | Cullin-4B | Q13620 |
| 340 | Ribosome-binding protein 1 | Q9P2E9 |
| 341 | Myristoylated alanine-rich C-kinase substrate | P29966 |
| 342 | DNA replication licensing factor MCM2 | P49736 |
| 343 | Ribosome biogenesis protein BMS1 homolog | Q14692 |
| 344 | E3 SUMO-protein ligase RanBP2 | P49792 |
| 345 | Isoform 3 of Transcriptional repressor p66-alpha | Q86YP4 |
| 346 | Structural maintenance of chromosomes protein 3 | Q9UQE7 |
| 347 | DNA mismatch repair protein Msh6 | P52701 |

| | | |
|---|---|---|
| 348 | Isoform 3 of Clathrin interactor 1 | Q14677 |
| 349 | Cytochrome b-c1 complex subunit 2, mitochondrial | P22695 |
| 350 | DnaJ homolog subfamily A member 1 | P31689 |
| 351 | Ribosomal RNA processing protein 1 homolog A | P56182 |
| 352 | Ubiquilin-1 | Q9UMX0 |
| 353 | Nuclear pore complex protein Nup93 | Q8N1F7 |
| 354 | Large ribosomal subunit protein uL29 | P42766 |
| 355 | Dolichyl-diphosphooligosaccharide--protein glycosyltransferase subunit 1 | P04843 |
| 356 | Serine/arginine-rich splicing factor 1 | Q07955 |
| 357 | U4/U6 small nuclear ribonucleoprotein Prp31 | Q8WWY3 |
| 358 | Isoform 2 of CAP-Gly domain-containing linker protein 1 | P30622 |
| 359 | Caprin-1 | Q14444 |
| 360 | 26S proteasome regulatory subunit 8 | P62195 |
| 361 | Hsp90 co-chaperone Cdc37 | Q16543 |
| 362 | Regulator of nonsense transcripts 3B | Q9BZI7 |
| 363 | Large ribosomal subunit protein bL12m | P52815 |
| 364 | GMP synthase [glutamine-hydrolyzing] | P49915 |
| 365 | Peroxisomal multifunctional enzyme type 2 | P51659 |
| 366 | Large ribosomal subunit protein P1 | P05386 |
| 367 | Small ribosomal subunit protein uS13 | P62269 |
| 368 | Ran GTPase-activating protein 1 | P46060 |
| 369 | 26S proteasome non-ATPase regulatory subunit 14 | O00487 |
| 370 | Peroxiredoxin-2 | P32119 |
| 371 | Keratin, type I cytoskeletal 18 | P05783 |
| 372 | Microtubule actin crosslinking factor 1 | H3BPE1 |
| 373 | Serine/arginine-rich splicing factor 3 | P84103 |
| 374 | Small ribosomal subunit protein eS27 | P42677 |
| 375 | HBS1-like protein | Q9Y450 |
| 376 | Proliferating cell nuclear antigen | P12004 |
| 377 | Lamin-B2 | Q03252 |
| 378 | 14-3-3 protein theta | P27348 |
| 379 | Phenylalanine--tRNA ligase alpha subunit | Q9Y285 |
| 380 | Histone-binding protein RBBP7 | Q16576 |
| 381 | Nucleolar protein 14 | P78316 |
| 382 | Eukaryotic translation initiation factor 3 subunit I | Q13347 |
| 383 | Peptidyl-prolyl cis-trans isomerase NIMA-interacting 1 | Q13526 |
| 384 | DnaJ homolog subfamily C member 9 | Q8WXX5 |
| 385 | Eukaryotic translation initiation factor 3 subunit B | P55884 |
| 386 | PDZ and LIM domain protein 1 | O00151 |
| 387 | Phosphoglycerate mutase 1 | P18669 |
| 388 | Elongation factor 1-beta | P24534 |
| 389 | Splicing factor 45 | Q96I25 |
| 390 | DNA ligase 3 | P49916 |
| 391 | Dynamin-1-like protein | O00429 |

| | | |
|---|---|---|
| 392 | Ras-related protein Rab-5C | P51148 |
| 393 | Lysine-rich nucleolar protein 1 | Q1ED39 |
| 394 | Protein phosphatase 1G | O15355 |
| 395 | WD repeat-containing protein 3 | Q9UNX4 |
| 396 | Dynactin subunit 1 | Q14203 |
| 397 | Regulation of nuclear pre-mRNA domain-containing protein 1B | Q9NQG5 |
| 398 | Kinesin-like protein KIF2C | Q99661 |
| 399 | Peptidyl-prolyl cis-trans isomerase-like 4 | Q8WUA2 |
| 400 | GTP-binding nuclear protein Ran | P62826 |
| 401 | Probable dimethyladenosine transferase | Q9UNQ2 |
| 402 | DNA-directed RNA polymerases I and III subunit RPAC1 | O15160 |
| 403 | Activator of 90 kDa heat shock protein ATPase homolog 1 | O95433 |
| 404 | Serine-threonine kinase receptor-associated protein | Q9Y3F4 |
| 405 | Zinc finger CCCH domain-containing protein 11A | O75152 |
| 406 | Coatomer subunit gamma-1 | Q9Y678 |
| 407 | Tubulin-specific chaperone A | O75347 |
| 408 | Voltage-dependent anion-selective channel protein 2 | P45880 |
| 409 | E3 ubiquitin-protein ligase RBBP6 | Q7Z6E9 |
| 410 | Parafibromin | Q6P1J9 |
| 411 | Isoform 2 of Replication factor C subunit 2 | P35250 |
| 412 | 26S proteasome regulatory subunit 4 | P62191 |
| 413 | Protein regulator of cytokinesis 1 | O43663 |
| 414 | Large ribosomal subunit protein uL18 | P46777 |
| 415 | Nucleolar protein 10 | Q9BSC4 |
| 416 | Nuclear receptor coactivator 5 | Q9HCD5 |
| 417 | RNA-binding protein 10 | P98175 |
| 418 | Structural maintenance of chromosomes protein 2 | O95347 |
| 419 | Pinin | Q9H307 |
| 420 | Programmed cell death protein 5 | O14737 |
| 421 | ATP-binding cassette sub-family F member 2 | Q9UG63 |
| 422 | Heat shock 70 kDa protein 4 | P34932 |
| 423 | Ataxin-2-like protein | Q8WWM7 |
| 424 | Succinate dehydrogenase [ubiquinone] flavoprotein subunit, mitochondrial | P31040 |
| 425 | Glutaredoxin-3 | O76003 |
| 426 | Small ribosomal subunit protein uS7 | P46782 |
| 427 | ATP synthase subunit gamma, mitochondrial | P36542 |
| 428 | Coiled-coil and C2 domain-containing protein 1A | Q6P1N0 |
| 429 | Small ribosomal subunit protein eS28 | P62857 |
| 430 | Ribonucleoside-diphosphate reductase subunit M2 | P31350 |
| 431 | Endophilin-A2 | Q99961 |
| 432 | Protein disulfide-isomerase A4 | P13667 |
| 433 | Small ribosomal subunit protein uS12 | P62266 |
| 434 | ESF1 homolog | Q9H501 |
| 435 | Protein SDA1 homolog | Q9NVU7 |

| | | |
|---|---|---|
| 436 | Metastasis-associated protein MTA2 | O94776 |
| 437 | Eukaryotic translation initiation factor 3 subunit A | Q14152 |
| 438 | Large ribosomal subunit protein eL19 | P84098 |
| 439 | Pre-mRNA-processing factor 40 homolog A | O75400 |
| 440 | Drebrin | Q16643 |
| 441 | G patch domain-containing protein 4 | Q5T3I0 |
| 442 | Bystin | Q13895 |
| 443 | Septin-9 | Q9UHD8 |
| 444 | Eukaryotic translation initiation factor 4B | P23588 |
| 445 | Nicotinamide phosphoribosyltransferase | P43490 |
| 446 | ATP-dependent 6-phosphofructokinase, liver type | P17858 |
| 447 | Vesicle-fusing ATPase | P46459 |
| 448 | Cleavage and polyadenylation specificity factor subunit 5 | O43809 |
| 449 | Eukaryotic translation initiation factor 6 | P56537 |
| 450 | ADP/ATP translocase 2 | P05141 |
| 451 | THO complex subunit 6 homolog | Q86W42 |
| 452 | Transcription elongation regulator 1 | O14776 |
| 453 | Exportin-5 | Q9HAV4 |
| 454 | Transportin-1 | Q92973 |
| 455 | Small ribosomal subunit protein uS15 | P62277 |
| 456 | DNA replication licensing factor MCM6 | Q14566 |
| 457 | Tubulin alpha-1C chain | Q9BQE3 |
| 458 | Nucleosome assembly protein 1-like 4 | Q99733 |
| 459 | Large ribosomal subunit protein uL22 | P18621 |
| 460 | Transcription elongation factor A protein 1 | P23193 |
| 461 | Large ribosomal subunit protein eL18 | Q07020 |
| 462 | 26S proteasome non-ATPase regulatory subunit 1 | Q99460 |
| 463 | U4/U6.U5 small nuclear ribonucleoprotein 27 kDa protein | Q8WVK2 |
| 464 | Heterogeneous nuclear ribonucleoprotein D-like | O14979 |
| 465 | Serine/arginine-rich splicing factor 9 | Q13242 |
| 466 | Filamin-B | O75369 |
| 467 | MICOS complex subunit MIC60 | Q16891 |
| 468 | Glutathione S-transferase P | P09211 |
| 469 | Isoform 2 of Eukaryotic peptide chain release factor GTP-binding subunit ERF3A | P15170 |
| 470 | Rab GDP dissociation inhibitor beta | P50395 |
| 471 | Large ribosomal subunit protein uL1 | P62906 |
| 472 | Spectrin beta chain, non-erythrocytic 1 | Q01082 |
| 473 | 3-hydroxyacyl-CoA dehydrogenase type-2 | Q99714 |
| 474 | Tubulin beta-2B chain | Q9BVA1 |
| 475 | Non-histone chromosomal protein HMG-14 | P05114 |
| 476 | Aldehyde dehydrogenase X, mitochondrial | P30837 |
| 477 | Isoform 4 of ADP-ribosylation factor-like protein 6-interacting protein 4 | Q66PJ3 |
| 478 | Splicing factor 3A subunit 3 | Q12874 |

| | | |
|---|---|---|
| 479 | Cyclin-dependent kinase 1 | P06493 |
| 480 | UDP-glucose 6-dehydrogenase | O60701 |
| 481 | Tryptophan--tRNA ligase, cytoplasmic | P23381 |
| 482 | Small ribosomal subunit protein uS2 | P08865 |
| 483 | U2 small nuclear ribonucleoprotein A' | P09661 |
| 484 | SUMO-activating enzyme subunit 1 | Q9UBE0 |
| 485 | Charged multivesicular body protein 4b | Q9H444 |
| 486 | Tubulin-folding cofactor B | Q99426 |
| 487 | GTPase-activating protein and VPS9 domain-containing protein 1 | Q14C86 |
| 488 | ATP-dependent RNA helicase | A0A8I5KSH6 |
| 489 | Non-histone chromosomal protein HMG-17 | P05204 |
| 490 | Isoform QKI7B of KH domain-containing RNA-binding protein QKI | Q96PU8 |
| 491 | Large ribosomal subunit protein eL30 | P62888 |
| 492 | Replication factor C subunit 3 | P40938 |
| 493 | Protein PRRC2A | P48634 |
| 494 | DNA mismatch repair protein Msh2 | P43246 |
| 495 | Leucine-rich repeat-containing protein 59 | Q96AG4 |
| 496 | RNA-binding protein FUS | P35637 |
| 497 | RNA-binding protein EWS | Q01844 |
| 498 | Eukaryotic translation initiation factor 5 | P55010 |
| 499 | Microfibrillar-associated protein 1 | P55081 |
| 500 | ATP-dependent RNA helicase DDX50 | Q9BQ39 |
| 501 | Eukaryotic translation initiation factor 4H | Q15056 |
| 502 | Pyruvate dehydrogenase E1 component subunit alpha, somatic form, mitochondrial | P08559 |
| 503 | Large ribosomal subunit protein eL14 | P50914 |
| 504 | DNA polymerase delta catalytic subunit | P28340 |
| 505 | DAZ-associated protein 1 | Q96EP5 |
| 506 | Ribosomal L1 domain-containing protein 1 | O76021 |
| 507 | WD repeat-containing protein 75 | Q8IWA0 |
| 508 | CUGBP Elav-like family member 1 | Q92879 |
| 509 | Targeting protein for Xklp2 | Q9ULW0 |
| 510 | Prelamin-A/C | P02545 |
| 511 | DDB1- and CUL4-associated factor 13 | Q9NV06 |
| 512 | Pre-mRNA 3'-end-processing factor FIP1 | Q6UN15 |
| 513 | Mitochondrial import inner membrane translocase subunit TIM44 | O43615 |
| 514 | Cellular tumor antigen p53 | A0A0U1RQC9 |
| 515 | cAMP-dependent protein kinase type II-alpha regulatory subunit | P13861 |
| 516 | Condensin complex subunit 1 | Q15021 |
| 517 | Smad nuclear-interacting protein 1 | Q8TAD8 |
| 518 | Bifunctional purine biosynthesis protein ATIC | P31939 |
| 519 | Small ribosomal subunit protein eS12 | P25398 |
| 520 | Importin-5 | O00410 |
| 521 | Isoform 2 of Neutral alpha-glucosidase AB | Q14697 |

| | | |
|---|---|---|
| 522 | Alpha-2-HS-glycoprotein | P02765 |
| 523 | p21-activated protein kinase-interacting protein 1 | Q9NWT1 |
| 524 | Nuclear pore complex protein Nup214 | P35658 |
| 525 | Cell division cycle and apoptosis regulator protein 1 | Q8IX12 |
| 526 | Basigin | P35613 |
| 527 | Heterogeneous nuclear ribonucleoprotein H2 | P55795 |
| 528 | Electron transfer flavoprotein subunit beta | P38117 |
| 529 | DNA fragmentation factor subunit alpha | O00273 |
| 530 | Small nuclear ribonucleoprotein Sm D2 | P62316 |
| 531 | Stathmin | P16949 |
| 532 | Membrane-associated progesterone receptor component 1 | O00264 |
| 533 | Adenylyl cyclase-associated protein 1 | Q01518 |
| 534 | Ras-related protein Rab-7a | P51149 |
| 535 | Isoform 3 of Nuclear pore complex protein Nup153 | P49790 |
| 536 | Small ribosomal subunit protein uS9 | P62249 |
| 537 | SWI/SNF complex subunit SMARCC1 | Q92922 |
| 538 | Catenin alpha-1 | P35221 |
| 539 | Translationally-controlled tumor protein | P13693 |
| 540 | Transmembrane emp24 domain-containing protein 9 | Q9BVK6 |
| 541 | Sorting nexin-2 | O60749 |
| 542 | GRB10 interacting GYF protein 2 | I1E4Y6 |
| 543 | Caldesmon | Q05682 |
| 544 | U4/U6 small nuclear ribonucleoprotein Prp3 | O43395 |
| 545 | Elongin-A | Q14241 |
| 546 | Rac GTPase-activating protein 1 | Q9H0H5 |
| 547 | 10 kDa heat shock protein, mitochondrial | P61604 |
| 548 | Protein LTV1 homolog | Q96GA3 |
| 549 | Survival of motor neuron-related-splicing factor 30 | O75940 |
| 550 | Small nuclear ribonucleoprotein Sm D3 | P62318 |
| 551 | Isoform 2 of Protein SGT1 homolog | Q9Y2Z0 |
| 552 | Nuclear migration protein nudC | Q9Y266 |
| 553 | U1 small nuclear ribonucleoprotein A | P09012 |
| 554 | Small ribosomal subunit protein uS19 | P62841 |
| 555 | Arginine and glutamate-rich protein 1 | Q9NWB6 |
| 556 | Ubiquitin carboxyl-terminal hydrolase 7 | Q93009 |
| 557 | Calpain small subunit 1 | P04632 |
| 558 | Small ribosomal subunit protein eS21 | P63220 |
| 559 | Myb-binding protein 1A | Q9BQG0 |
| 560 | tRNA methyltransferase 10 homolog C | Q7L0Y3 |
| 561 | Succinate--CoA ligase [ADP/GDP-forming] subunit alpha, mitochondrial | P53597 |
| 562 | Transforming protein RhoA | P61586 |
| 563 | Adenosylhomocysteinase | P23526 |
| 564 | Serine/arginine repetitive matrix protein 1 | Q8IYB3 |
| 565 | Stomatin-like protein 2, mitochondrial | Q9UJZ1 |

| | | |
|---|---|---|
| 566 | tRNA (guanine(26)-N(2))-dimethyltransferase | Q9NXH9 |
| 567 | Splicing factor U2AF 35 kDa subunit | Q01081 |
| 568 | Destrin | P60981 |
| 569 | Large ribosomal subunit protein eL15 | P61313 |
| 570 | V-type proton ATPase subunit B, brain isoform | P21281 |
| 571 | Endophilin-B1 | Q9Y371 |
| 572 | Nucleoporin 155 | E9PF10 |
| 573 | Eukaryotic translation initiation factor 3 subunit E | A0A7I2V3S3 |
| 574 | PEST proteolytic signal-containing nuclear protein | Q8WW12 |
| 575 | Phenylalanine--tRNA ligase beta subunit | Q9NSD9 |
| 576 | Histone H2AX | P16104 |
| 577 | Malate dehydrogenase, cytoplasmic | P40925 |
| 578 | Glutamate dehydrogenase 1, mitochondrial | P00367 |
| 579 | RRM domain-containing protein (Fragment) | A0A590UK80 |
| 580 | Uncharacterized protein C11orf98 | E9PRG8 |
| 581 | Small nuclear ribonucleoprotein Sm D1 | P62314 |
| 582 | Zinc finger CCCH domain-containing protein 15 | Q8WU90 |
| 583 | Cytosolic acyl coenzyme A thioester hydrolase | O00154 |
| 584 | Protein SETSIP | P0DME0 |
| 585 | Large ribosomal subunit protein uL30 | P18124 |
| 586 | Charged multivesicular body protein 2b | Q9UQN3 |
| 587 | N-alpha-acetyltransferase 15, NatA auxiliary subunit | Q9BXJ9 |
| 588 | Isoform 2 of Ubiquitin carboxyl-terminal hydrolase isozyme L5 | Q9Y5K5 |
| 589 | Alanine--tRNA ligase, mitochondrial | Q5JTZ9 |
| 590 | Pumilio homolog 1 | Q14671 |
| 591 | Histone-binding protein RBBP4 | Q09028 |
| 592 | Large ribosomal subunit protein eL21 | P46778 |
| 593 | Protein FAM50A | Q14320 |
| 594 | Large ribosomal subunit protein uL2 | P62917 |
| 595 | Regulation of nuclear pre-mRNA domain-containing protein 1A | Q96P16 |
| 596 | ATP synthase subunit O, mitochondrial | A0A494C0K9 |
| 597 | Serine/threonine-protein phosphatase 1 regulatory subunit 10 | Q96QC0 |
| 598 | Glucosidase 2 subunit beta | P14314 |
| 599 | Small ribosomal subunit protein uS4 | P46781 |
| 600 | Ribosome biogenesis protein WDR12 | Q9GZL7 |
| 601 | Asparagine--tRNA ligase, cytoplasmic | O43776 |
| 602 | Voltage-dependent anion-selective channel protein 1 | P21796 |
| 603 | Endoplasmic reticulum resident protein 29 | P30040 |
| 604 | Chromobox protein homolog 5 | P45973 |
| 605 | Large ribosomal subunit protein uL15 | P46776 |
| 606 | Putative methyltransferase C9orf114 | Q5T280 |
| 607 | Dynactin subunit 2 | Q13561 |
| 608 | Calmodulin-1 | P0DP23 |
| 609 | Isoform 1 of Four and a half LIM domains protein 1 | Q13642 |
| 610 | Myotrophin | P58546 |

| | | |
|---|---|---|
| 611 | Histone deacetylase complex subunit SAP18 | O00422 |
| 612 | Ezrin | P15311 |
| 613 | Nucleolar protein 12 | Q9UGY1 |
| 614 | Telomere-associated protein RIF1 | Q5UIP0 |
| 615 | U4/U6 small nuclear ribonucleoprotein Prp4 | O43172 |
| 616 | Thioredoxin domain-containing protein 5 | Q8NBS9 |
| 617 | Dual specificity mitogen-activated protein kinase kinase 1 | Q02750 |
| 618 | DnaJ homolog subfamily C member 8 | O75937 |
| 619 | High mobility group protein B2 | P26583 |
| 620 | AMP deaminase 2 | Q01433 |
| 621 | NF-kappa-B-activating protein | Q8N5F7 |
| 622 | NEDD8-conjugating enzyme Ubc12 | P61081 |
| 623 | Large ribosomal subunit protein uL13 | P40429 |
| 624 | Ubiquitin carboxyl-terminal hydrolase isozyme L1 | P09936 |
| 625 | Lysine-specific demethylase RSBN1L | Q6PCB5 |
| 626 | Small nuclear ribonucleoprotein-associated proteins B and B' | P14678 |
| 627 | Large ribosomal subunit protein eL28 | P46779 |
| 628 | C-terminal-binding protein 1 | Q13363 |
| 629 | Large ribosomal subunit protein eL20 | Q02543 |
| 630 | Small ribosomal subunit protein eS25 | P62851 |
| 631 | Coiled-coil domain-containing protein 124 | Q96CT7 |
| 632 | WD repeat-containing protein 43 | Q15061 |
| 633 | Transcription factor A, mitochondrial | Q00059 |
| 634 | Isoform 2 of Jupiter microtubule associated homolog 1 | Q9UK76 |
| 635 | Large ribosomal subunit protein eL34 | P49207 |
| 636 | Protein arginine N-methyltransferase 5 | O14744 |
| 637 | Isoform 2 of B-cell receptor-associated protein 31 | P51572 |
| 638 | Serine/arginine-rich splicing factor 2 | Q01130 |
| 639 | Hemoglobin subunit alpha | P69905 |
| 640 | eIF5-mimic protein 2 | Q7L1Q6 |
| 641 | Trifunctional enzyme subunit beta, mitochondrial | P55084 |
